# Assembly and levels of P-TEFb depend on reversible phosphorylation of cyclin T1

**DOI:** 10.1101/2021.05.25.445695

**Authors:** Fang Huang, Trang T. T. Nguyen, Ignacia Echeverria, Rakesh Ramachandran, Daniele C. Cary, Hana Paculova, Andrej Sali, Arthur Weiss, B. Matija Peterlin, Koh Fujinaga

## Abstract

The positive transcription elongation factor b (P-TEFb) is a critical co-activator for transcription of most cellular and viral genes, including those of HIV. While P-TEFb is regulated by 7SK snRNA in proliferating cells, it is absent in quiescent and terminally differentiated cells, which has remained unexplored. In these cells, we found that CycT1 not bound to CDK9 is rapidly degraded. Moreover, productive CycT1:CDK9 interactions require phosphorylation of two threonine residues (Thr143 and Thr149) in CycT1 by PKC. Conversely, PP1 dephosphorylates these sites. Thus, PKC inhibitors or removal of PKC by chronic activation results in P-TEFb disassembly and CycT1 degradation. This finding not only recapitulates P-TEFb depletion in resting CD4+ T cells but also in anergic T cells. Importantly, our studies reveal mechanisms of P-TEFb inactivation underlying T cell quiescence, anergy, and exhaustion as well as proviral latency and terminal differentiation of cells.

## Introduction

In eukaryotic cells, coding gene expression starts with transcription of DNA to RNA by RNA polymerase II (RNAPII) in the nucleus. This process consists of initiation, promoter clearance, capping, elongation and termination, upon which the transcribed single strand RNA is cleaved and poly-adenylated before transport to the cytoplasm (Lis, 2019; Peterlin & Price, 2006; Proudfoot, 2016; Zhou, Li, & Price, 2012). Of interest, promoters of most inactive and inducible genes are already engaged by stalled RNAP II, which pauses after transcribing 20-100 nucleotides long transcripts (Rahl et al., 2010). The pause of RNAP II is caused by two factors, the negative elongation factor (NELF) (Yamaguchi et al., 1999) and DRB sensitivity inducing factor (DSIF) (Wada, Takagi, Yamaguchi, Ferdous, et al., 1998). Clearance of RNAP II from promoters requires the phosphorylation of its C-terminal domain (CTD) at position 5 (Ser5) in the tandemly repeated heptapeptide (52 repeats of Tyr1-Ser2-Pro3-Thr4-Ser5-Pro6-Ser7) by cyclin-dependent kinase 7 (CDK7) from the transcription factor-II H (TFIIH) (Chapman, Heidemann, Hintermair, & Eick, 2008). The release of RNAP II for productive elongation requires the positive transcription factor b (P-TEFb), which is composed of cyclin-dependent kinase 9 (CDK9) and C-type cyclins T1 or T2 (CycT1 or CycT2) (Zhou et al., 2012). Compared to the largely restricted expression of CycT2, CycT1 is expressed ubiquitously (Peng, Zhu, Milton, & Price, 1998). N-terminus of CycT1 contains two highly conserved cyclin boxes for CDK9 binding, followed by the Tat-TAR recognition motif (TRM), a coil-coiled motif, the histidine (His) rich motif that binds to the CTD and a C-terminal PEST motif (Taube, Lin, Irwin, Fujinaga, & Peterlin, 2002; Wei, Garber, Fang, Fischer, & Jones, 1998). CDK9 is a Ser/Thr proline-directed kinase (i.e. PITALRE) (Grana et al., 1994) that phosphorylates Spt5 in DRB and NELF-E, that relieve the pausing of RNAPII (Fujinaga et al., 2004; Ivanov, Kwak, Guo, & Gaynor, 2000). After phosphorylation, NELF is released and DSIF is converted to an elongation factor (Wada, Takagi, Yamaguchi, Watanabe, & Handa, 1998). Serine at position 2 (Ser2) in the CTD is also phosphorylated by CDK9 before RNAPII’s transition to productive elongation (Peterlin & Price, 2006). Thus, P-TEFb is a critical factor for transcriptional elongation and co-transcriptional processing by RNAPII.

In the organism, the kinase activity of P-TEFb is kept under a tight control to maintain the appropriate state of growth and proliferation of cells. To ensure this balance, a large complex, known as the 7SK small nuclear ribonucleoprotein (7SK snRNP) sequesters a large amount of P-TEFb (from 50% to 90% in different cells) in an inactive state (Peterlin, Brogie, & Price, 2012). 7SK snRNP consists of the abundant 7SK small nuclear RNA (7SK snRNA), hexamethylene bisacetamide (HMBA)-inducible mRNAs 1 and 2 (HEXIM1/2) proteins, La-related protein 7 (LARP7) and methyl phosphate capping enzyme (MePCE) (Michels & Bensaude, 2008; Zhou et al., 2012). Proper levels and activities of P-TEFb ensure appropriate responses to external stimuli. They also maintain states of differentiation, growth and proliferation of cells (AJ, Bugai, & Barboric, 2016; Fujinaga, 2020). Dys-regulation of the P-TEFb equilibrium contributes to various diseases such as solid tumors (mutation in LARP7) leukemias and lymphomas (DNA translocations leading to aberrant recruitment of P-TEFb), and cardiac hypertrophy (inactivation of HEXIM1) (Franco, Morales, Boffo, & Giordano, 2018). Cellular stresses such as ultraviolet (UV) irradiation and heat, as well as various small compound such as histone deacetylase inhibitors (HDACi), HMBA and bromodomain extra-terminal domain (BET) inhibitors (JQ1), also promote the release of P-TEFb from the 7SK snRNP in a reversible manner (Zhou et al., 2012). Despite these important cellular responses, different viruses, such as HIV, HTLV, EBV, HSV, HCMV and others, have evolved different strategies to utilize P-TEFb for their own replication (Zaborowska, Isa, & Murphy, 2016). For example, the HIV transactivator of transcription (Tat) not only binds to free P-TEFb but also promotes the release of P-TEFb from 7SK snRNP. Tat:P-TEFb then binds to the trans-activation response (TAR) RNA stem-loop to activate the transcription of viral genes (Selby & Peterlin, 1990; Wei et al., 1998). Additionally, the establishment of viral latency in quiescent cells parallels the disappearance of P-TEFb, which can be reversed by cell activation (Rice, 2019).

In activated and proliferating cells, high levels of P-TEFb are found. In contrast, they are vanishingly low in resting cells, especially monocytes and memory T cells. While levels of CDK9 persist, those of CycT1 are greatly reduced. At the same time, transcripts for CycT1 and CDK9 remain high in all these cells (Garriga et al., 1998; Ghose, Liou, Herrmann, & Rice, 2001). Based on existing studies, one can conclude that P-TEFb falls apart when cells become quiescent. In these cells, CDK9 is stabilized by chaperone proteins HSP70 and HSP90 (O’Keeffe, Fong, Chen, Zhou, & Zhou, 2000).

For CycT1, it was thought that its translation is inhibited by RNAi (Chiang & Rice, 2012; Sung & Rice, 2009). In contrast, we find that CycT1 is rapidly degraded in these cells. We also identified post-translational modifications that lead to the assembly and disassembly of P-TEFb, which involves specific kinases and phosphatases. Importantly, the unbound CycT1 protein can be stabilized by proteasomal inhibitors. This situation appears uncannily reminiscent of cell cycle CDKs that are also regulated by similar post-translational mechanisms.

## Results

### Critical residues in CycT1 (Thr143 and Thr149) are required for its binding to CDK9

Previously, residues in the N-terminal region of CycT1 (positions 1-280, CycT1(280), including cyclin boxes, positions 30-248) (Fig. 1A) were found to be required for interactions between CycT1 and CDK9 (Garber et al., 1998). In particular, a substitution of the leucine to proline at position 203 (L203P) or four substitutions from a glutamic to aspartic acid at position 137 and threonine to alanine at positions 143, 149 and 155 (4MUT) completely abolished this binding (Kuzmina et al., 2014) (Fig. 1A). We created a similar set of mutant CycT1 proteins in the context of the full length CycT1 and truncated CycT1(280) proteins (Fig. 1A). Next, we defined further critical residues involved in CycT1:CDK9 interactions, especially the three adjacent threonine residues in the cyclin box (CycT1T3A, Fig. 1A). First, these mutant CycT1 proteins were expressed in 293T cells. Next, interactions between mutant CycT1 proteins and the endogenous CDK9 protein were analyzed by co-immunoprecipitation (co-IP) (Figs. 1B-1D).

**Fig. 1.**
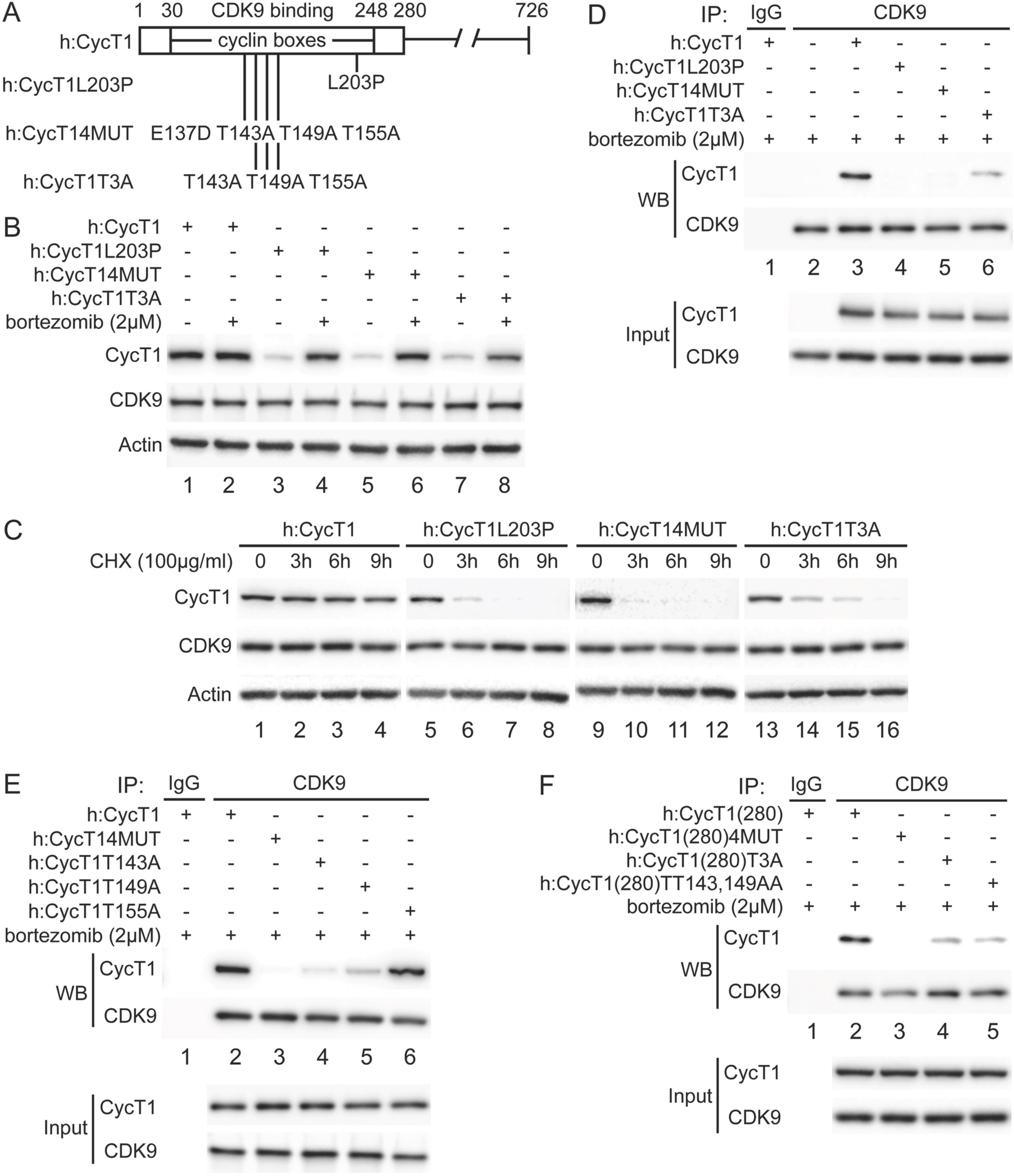
Critical residues in CycT1 (Thr143 and Thr149) are required for its binding to CDK9. A. Diagram of CycT1 and indicated mutant CycT1 proteins. The full length human CycT1 protein contains 726 residues. Two cyclin boxes are found between positions 30 and 248. Critical residues for CDK9 binding include Thr143, Thr149 and Thr155. Glu137 and Leu203 flank these sites. Presented are critical mutations in CycT1 that form the basis of this study. B. Mutant CycT1 proteins are unstable. CycT1 and three indicated mutant CycT1 proteins were expressed in 293T cells, which were untreated (lances 1, 3, 5 and 7) or treated with 2µM bortezomib for 12 h (lanes 2, 4, 6 and 8) before cell lysis. Levels of CycT1 (1st panel), CDK9 (2nd panel), and the loading control actin (3rd panel) proteins were detected with anti-HA, anti-CDK9, and anti-β-actin antibodies, respectively, by WB. Gels are marked as follows: IP, IPed proteins, above the panels; next, presence and absence of co-IPed proteins is denoted by (+) and (-) signs; same for the inclusion and concentration of bortezomib; WB, western blot of co-IPed proteins; Input, western blot of input proteins. C. Mutant CycT1 proteins are unstable. CycT1 and three indicated mutant CycT1 proteins were expressed in 293T cells, which were untreated (lanes 1, 5, 9 and 13) or treated with 100 µg/ml cycloheximide (CHX) for 3-9 h (lanes 2 to 4; 6 to 8; 10 to 12; 14 to 16) before cell lysis. Levels of CycT1 (1st panel), CDK9 (2nd panel), and the loading control actin (3rd panel) proteins were detected with anti-HA, anti-CDK9, and anti-β-actin antibodies, respectively, by WB. D. Interactions between CDK9 and mutant CycT1 proteins are impaired. In 293T cells treated with 2µM bortezomib, co-IPs with CDK9 are presented in top two panels. 3rd and 4th panels contain input levels of CycT1 and CDK9 proteins. E. Interactions between CDK9 and point mutant CycT1 proteins are impaired. In 293T cells treated with 2µM bortezomib, co-IPs with CDK9 are presented in top two panels. 3rd and 4th panels contain input levels of CycT1 and CDK9 proteins. F. Interactions between CDK9 and truncated mutant CycT1(280) proteins are impaired. In 293T cells treated with 2µM bortezomib, co-IPs with CDK9 are presented in top two panels. 3rd and 4th panels contain input levels of CycT1(280) and CDK9.

Mutant CycT1 proteins were poorly expressed in 293T cells (Fig. 1B, 1st panel, lanes 3, 5 and 7). Expression levels of these proteins were restored by incubating these cells with the potent and clinically approved proteasomal inhibitor bortezomib (Fig.1B, 1st panel, compare lanes 4, 6 and 8 to lanes 3, 5 and 7), indicating that these mutant CycT1 proteins are highly unstable in cells. To confirm this finding, the half life of the mutant CycT1 proteins was measured by cycloheximide (CHX) pulse-chase experiments (Fig. 1C). Whereas levels of the wild type CycT1 protein remained unchanged after cycloheximide treatment (Fig. 1C, 1st panel, lanes 1 to 4), those of three mutant CycT1 proteins (CycT1L203P, CycT14MUT and CycT1T3A) decreased rapidly with the half life of ∼3 h, ∼2.5 h, and ∼6 h, respectively (Fig. 1C, 1st panel, lanes 5 to 8; lanes 9 to 12; lanes 13 to 16). Moreover, levels of endogenous CDK9 protein were not changed under bortezomib or cycloheximide treatment (Fig. 1A and 1B, 2nd panels). When protein levels were restored by bortezomib, mutant CycT1 proteins (CycT1L203P and CycT14MUT) did not interact with CDK9 (Fig. 1D, 1st panel, compare lanes 4 and 5 to lane 3). Similarly, interactions between the mutant CycT1T3A protein and CDK9 were significantly decreased (Fig. 1D, 1st panel, compare lane 6 to lane 3, ∼7.8-fold reduction). We further examined whether all these threonine residues are important for binding to CDK9 (Fig. 1E). Thus, mutant CycT1 proteins with single threonine substitution were created and examined for CycT1:CDK9 interactions. As presented in Fig. 1E, whereas mutant CycT1T143A and CycT1T149A proteins exhibited significantly impaired binding to CDK9, the mutant CycT1T155A protein did not (1st panel, compare lanes 4 and 5 to lanes 2 and 6, ∼5.2-fold and ∼3.7-fold reduction).

These results indicate that Thr143 and Thr149, but not Thr155, are critical residues in CycT1 for binding to CDK9. Finally, the mutant CycT1(280) protein with double threonine substitution to alanine (CycT1(280)TT143,149AA) demonstrated reduced binding to CDK9 to a similar extent as the mutant CycT1(280)T3A protein (Fig.1F, 1st panel, compare lanes 4 and 5 to lane 2, ∼7.2-fold vs ∼8.1-fold reduction). Taken together, we conclude that two threonine residues (Thr143 and Thr149) in CycT1 are critical for its binding to CDK9 to form P-TEFb, and that mutant CycT1 proteins that do not interact or poorly interact with CDK9 are rapidly degraded by the proteasome.

### Phosphorylation of Thr143 and Thr149 in CycT1 contributes to its binding to CDK9

Since threonine residues are potential phosphorylation sites, we examined whether phosphorylation of Thr143 and/or Thr149 in CycT1 contributes to P-TEFb assembly. 293T cells ectopically expressing CycT1 (Fig. 2A) or CycT1(280) (Fig. 2B) were treated with the potent protein phosphatase inhibitor okadaic acid for 1.5 h prior to cell lysis.

**Fig. 2.**
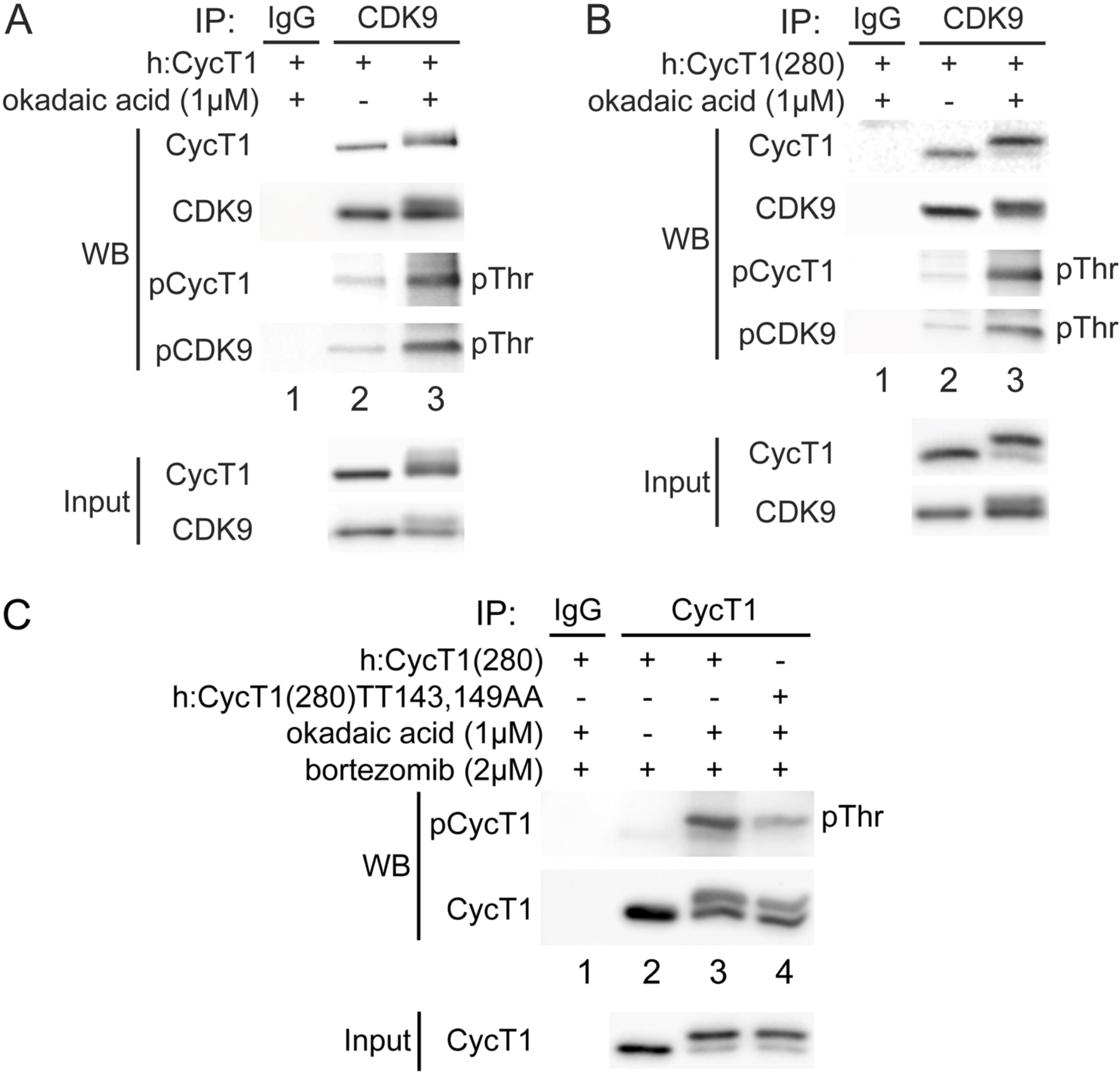
Phosphorylation of Thr143 and Thr149 in CycT1 contributes to its binding to CDK9. A. Threonine phosphorylation is detected in the full length CycT1 protein. CycT1 and CDK9 proteins were co-expressed in the presence and absence of 1µM okadaic acid (+/− signs on top) in 293T cells. Co-IPs with CDK9 were then probed with anti-phospho-threonine (pThr) antibodies in the middle two panels. 5th and 6th panels contain input levels of CycT1 and CDK9 proteins. B. Threonine phosphorylation is detected in CycT1(280). CycT1(280) and CDK9 were co-expressed in the presence and absence of 1µM okadaic acid (+/− signs on top) in 293T cells. Co-IPs with CDK9 were then probed with anti-pThr antibodies in the middle two panels. 5th and 6th panels contain input levels of CycT1 and CDK9 proteins. C. Thr143 and Thr149 are major phospho-threonine residues in CycT1. CycT1(280) or the mutant CycT1(280)TT143,149AA protein was expressed in the presence of 2µM bortezomib as well as in the presence or absence of 1µM okadaic acid (+/− signs on top) in 293T cells. IPs with CycT1 were then probed with anti-pThr antibodies in the top panel. 4th and 5th panels contain input levels of CycT1 and CDK9.

Following the co-IP of CDK9, its phosphorylation was analyzed with anti-phospho-threonine (pThr) antibodies by western blotting (WB). Phospho-threonine signals were increased in both CycT1 and CycT1(280) proteins in the presence of a high concentration of okadaic acid, which inhibits serine/threonine protein phosphatases 1 (PP1) (Fig. 2A and 2B, 3rd panels, lanes 3), but not by a low concentration of okadaic acid (data not presented), which inhibits PP2A. Confirming these inhibitory effects by okadaic acid, CycT1 and CDK9 bands shifted upwards by this treatment, and only these upper bands were detected with anti-pThr antibodies (Fig. 2). Furthermore, interactions between CycT1 and CDK9 were increased by the high concentration of okadaic acid treatment (Fig. 2A, 1st panel, compare lane 3 to lane 2, ∼5.1-fold increase). In addition, CycT1 co-IPed with CDK9 was heavily phosphorylated at threonine residues (Fig. 2A, 3rd panel, compare lane 3 to lane 2, ∼19-fold increase). Under the same conditions, CDK9 was also heavily phosphorylated (Fig. 2A, 4th panel, compare lane 3 to lane 2, ∼8.9-fold increase). Similarly, interactions between CycT1(280) and CDK9 were also increased by okadaic acid (Fig. 2B, 1st panel, compare lane 3 to lane 2, ∼4.5-fold increase). Threonine phosphorylation of CDK9-associated CycT1(280) was also increased (Fig. 2B, 3rd panel, compare lane 3 to lane 2, ∼21-fold increase). Increased phosphorylated CDK9 was also detected (Fig. 2B, 4th panel, compare lane 3 to lane 2, ∼9.4-fold increase).

An online database for phosphorylation site prediction (NetPhos 3.1, developed by Technical University of Denmark) scores Thr143 and Thr149 above the threshold value (default 0.5), indicating that these threonines are potential phosphorylation sites (Fig. S1).To verify that Thr143 and Thr149 are the main phosphorylation sites in CycT1, levels of total threonine phosphorylation were compared between CycT1(280) and mutant CycT1(280)TT143,149AA proteins in the presence of high concentration of okadaic acid. As presented in Fig. 2C, levels of threonine phosphorylation were significantly reduced in the mutant CycT1(280)TT143,149AA protein compared to CycT1(280) (1st panel, compare lane 4 to lane 3, ∼5.2-fold reduction), indicating that Thr143 and Thr149 are major threonine phosphorylation sites. Taken together, we conclude that the phosphorylation of Thr143 and Thr149 in CycT1 is essential for its binding to CDK9, and that PP1 is involved in the dephosphorylation of CycT1.

### Residues in CycT1 and CDK9 that regulate the assembly of P-TEFb

These two threonine residues (Thr143 and Thr149) are located in cyclin boxes that are required for CDK9 binding. To understand further the exact roles of these two sites in the interface between CycT1 and CDK9, we analyzed molecular interactions between specific side chain residues in these proteins based on the crystal structure of the human P-TEFb complex (PDB access code 3MI9) (Baumli et al., 2008; Tahirov et al., 2010). Further, we modeled phosphorylated Thr143 and Thr149 using this crystal structure. Phosphates were added to Thr143 and Thr149 of the 3MI9 crystal structure, followed by energy minimization using the UCSF Chimera program (Pettersen et al., 2004). The three dimensional representation was created using Pymol (PyMOL Molecular Graphics System, Version 2.0, Schrödinger, LLC.) (Fig. 3A).

**Fig.3.**
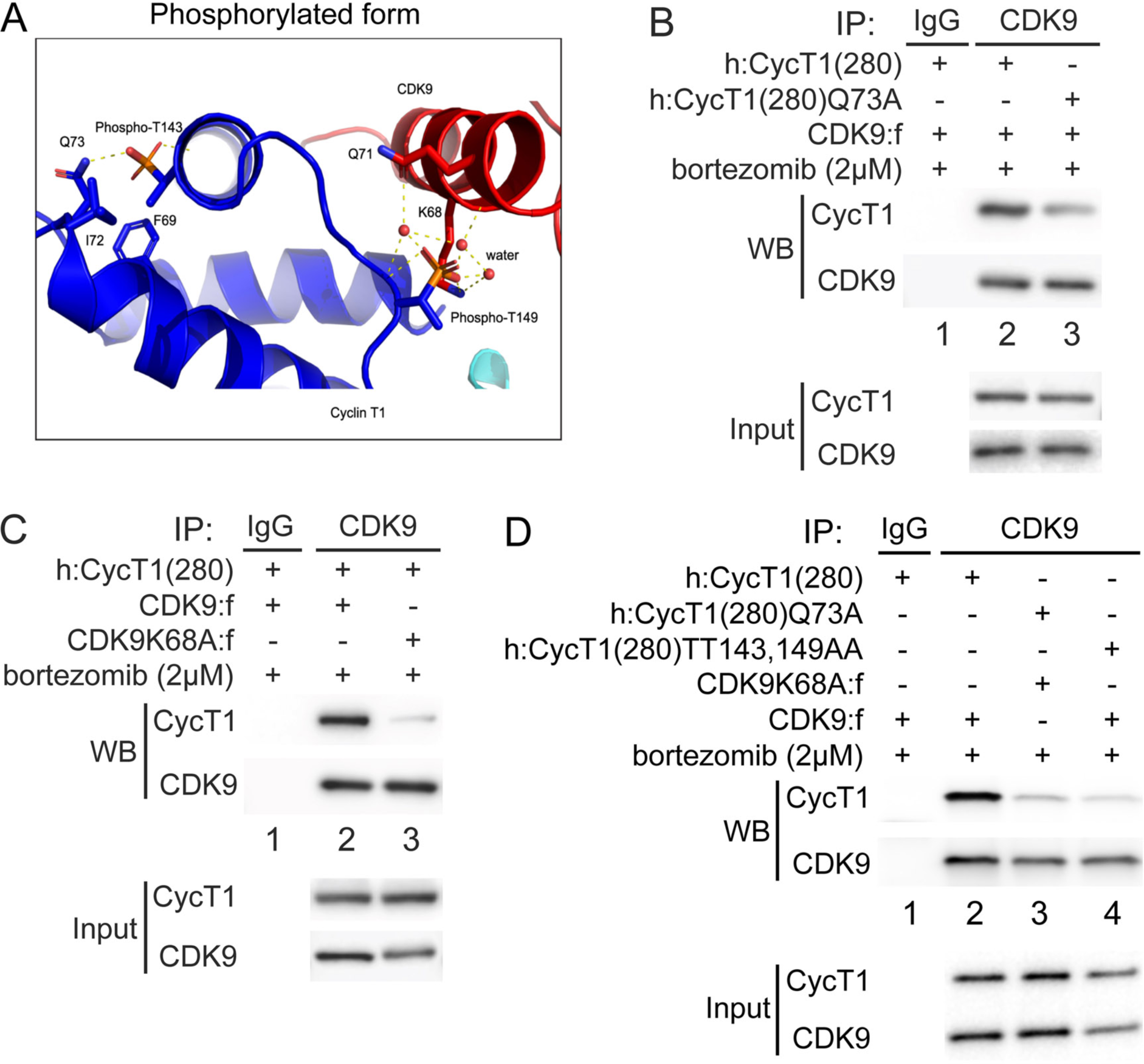
Phosphorylation of Thr143 and Thr149 stabilizes the interface between CycT1 and CDK9. A. Model of the complex phosphorylated at Thr143 and Thr149 in CycT1. Residues interacting with Thr143 and Thr149 are Gln73 in CycT1 and Lys68 in CDK9, respectively. Dashed lines represent interactions with water molecules. The model was obtained by adding phosphates to Thr143 and Thr149 in the crystal structure (PDB ID 3MI9), followed by energy minimization to resolve steric clashes. B. Gln73 is targeted by phosphorylated Thr143 in CycT1. CycT1(280) or the mutant CycT1(280)Q73A protein and CDK9 were co-expressed in the presence of 2µM bortezomib (+/− signs on top) in 293T cells. Co-IPs with CDK9 are presented in top panels (WB). 3rd and 4th panels contain input levels of CycT1 and CDK9 (input). C. Lys68 in CDK9 is targeted by phosphorylated Thr149 in CycT1. CDK9 or mutant CDK9K68A protein and CycT1(280) were co-expressed in the presence of 2µM bortezomib (+/− signs on top) in 293T cells. Co-IPs with CDK9 are presented in top panels (WB). 3rd and 4th panels contain input levels of CycT1 and CDK9 (input). D. Mutations of K68A in CDK9 and Q73A in CycT1 attenuate cooperatively the binding between CycT1 and CDK9, equivalently to the mutant CycT1TT143,149AA protein. CycT1(280) or the mutant CycT1(280)Q73A protein and CDK9 or the mutant CDK9K68A protein were co-expressed in the presence of 2µM bortezomib (+/− signs on top) in 293T cells. Co-IPs with CDK9 are presented in top panels (WB). 3rd and 4th panels contain input levels of CycT1 and CDK9 (input).

As presented in Fig. 3A, the structure of the unphosphorylated form can readily accommodate the phosphorylation of Thr143 and Thr149 in CycT1, potentially stabilizing intramolecular interactions with Gln73 in CycT1 and intermolecular interactions with Lys68 in CDK9, respectively. The role of these predicted sites was confirmed experimentally by co-IPs using mutant proteins. CDK9 or the mutant CDK9K68A protein were co-expressed with CycT1(280) or the mutant CycT1(280)Q73A proteins in the presence of bortezomib for 12 h before co-IP. As presented in Fig. 3B, compared to CycT1(280), the mutant CycT1(280)Q73A protein exhibited a lower affinity for CDK9 (1st panel, compare lane 3 to lane 2, ∼4.5-fold reduction). Similarly, compared to CDK9, the mutant CDK9K68A protein exhibited a lower affinity for CycT1(280) (Fig. 3C, 1st panel, compare lane 3 to lane 2, ∼5.9-fold reduction). Finally, interactions between the mutant CycT1(280)Q73A and CDK9K68A proteins were reduced to a similar extent to those between the mutant CycT1(280)TT143,149AA and CDK9 proteins, compared to the positive control with CDK9 and CycT1(280) (Fig. 3D, 1st panel, compare lanes 3 and 4 to lane 2, ∼7.5-fold vs ∼8-fold reduction, respectively). Taken together, phosphates on Thr143 and Thr149 in CycT1 are essential for the assembly and stability of P-TEFb.

### PKC inhibitors impair interactions between CycT1 and CDK9 and promote CycT1 degradation

In resting and memory T cells, levels of CycT1 are vanishingly low. In previous sections, we discovered that Thr143 and Thr149 are phosphorylated in the stable P-TEFb complex. These residues can be dephosphorylated by PP1. Kinases that cause this phosphorylation remained unknown. Nevertheless, using kinase prediction programs (NetPhos 3.1), these residues lie in separate PKC consensus sites. While Thr143 received the highest score, Thr149 could also be a target for PKC or other kinases.

These findings implied that PKC family members are kinases that phosphorylate Thr143 and/or Thr149. To confirm that PKC promotes the phosphorylation and the stability of CycT1, several PKC inhibitors were introduced to different cells (Fig. 4). Of these, staurosporine exhibited the most significant inhibition in 293T cells. As presented in Fig. 1B and 1C, the exogenous CycT1 protein is very stable in cells. Next, increasing amount of staurosporine were added to cells 12 h prior to cell lysis. Staurosporine reduced levels of CycT1 in these cells in a dose dependent manner (Fig. S2A, 1st panel, compare lanes 2 and 3 to lane 1, ∼4-fold and ∼16-fold reduction, respectively).

**Fig. 4.**
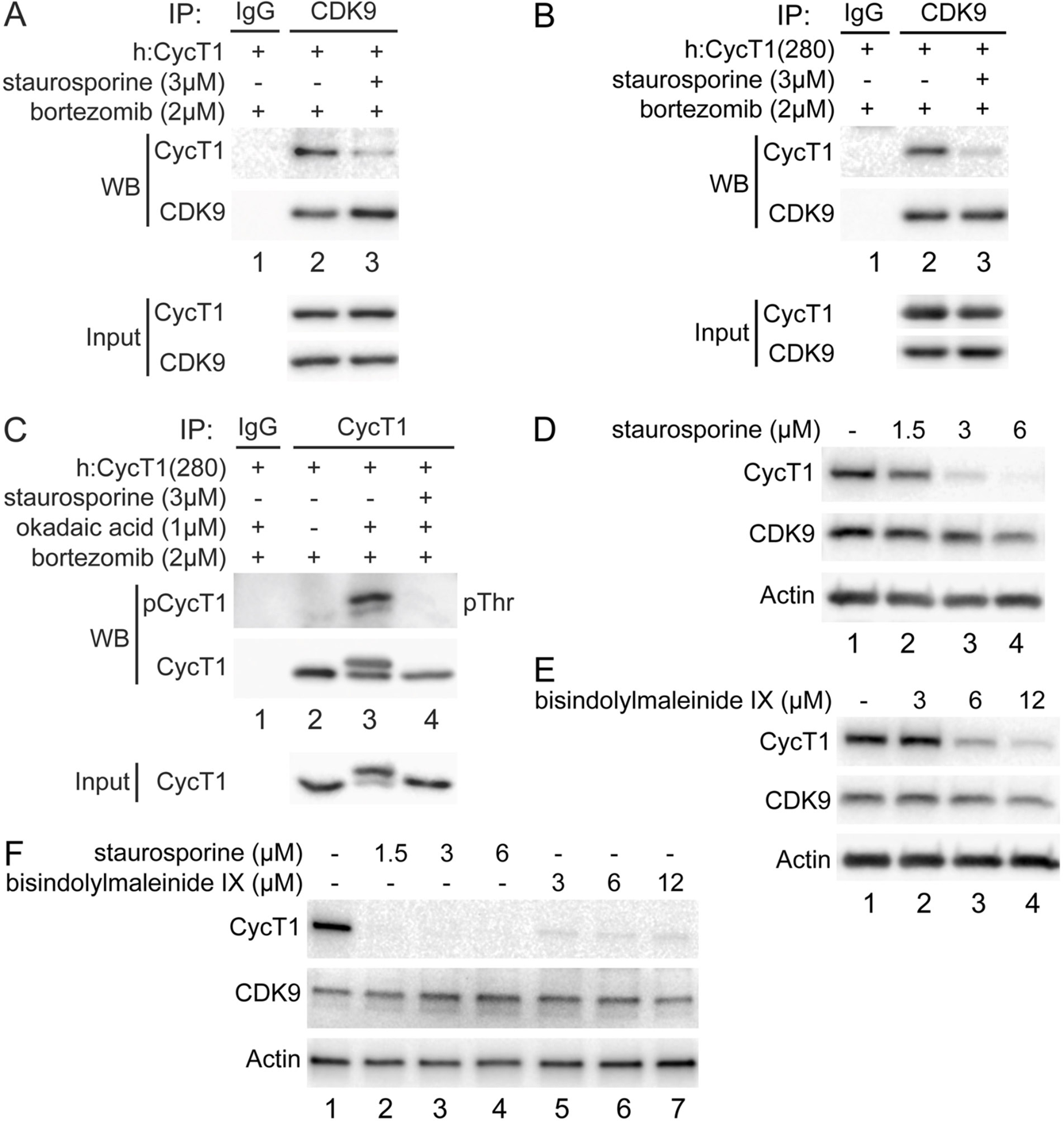
PKC inhibitors impair interactions between CycT1 and CDK9, and promote CycT1 degradation. A. PKC inhibitors impair interactions between CycT1 and CDK9. CycT1 and CDK9 were co-expressed in the presence or absence of 3µM staurosporine and 2µM bortezomib (+/− signs on top) in 293T cells. Co-IPs with CDK9 are presented in the top panels (WB). 3rd and 4th panels contain input levels of CycT1 and CDK9 (input). B. PKC inhibitors impair interactions between CycT1(1-280) and CDK9. CycT1(280) and CDK9 were co-expressed in the presence or absence of 3µM staurosporine and 2µM bortezomib (+/− signs on top) in 293T cells. Co-IPs with CDK9 are presented in the top panels (WB). 3rd and 4th panels contain input levels of CycT1(280) and CDK9 (input). C. PKC inhibitor staurosporine inhibits threonine phosphorylation of CycT1. CycT1(280) was expressed in the presence or absence of 3µM staurosporine, 1µM okadaic acid and 2µM bortezomib (+/− signs on top) in 293T cells. IPs with CycT1 are presented in the top panels (WB). Phosphorylated proteins were visualized with anti-pThr antibodies (top panel). 3rd panel contains input levels of CycT1(280) (Input). D. Staurosporine decreases CycT1 levels in a dose dependent manner. Jurkat cells were untreated (lane 1) or treated with increasing doses of staurosporine (1.5µM, 3µM and 6µM) (lanes 2 to 4) for 12 h before cell lysis. Levels of CycT1 (1st panel), CDK9 (2nd panel) and the loading control actin (3rd panel) proteins were detected with anti-CycT1, anti-CDK9 and anti-β-actin antibodies, respectively, by WB. E. PKC inhibitor bisindolylmaleinide IX decreases CycT1 levels in a dose dependent manner. Jurkat cells were untreated (lane 1) or treated with increasing doses of bisindolylmaleinide IX (3µM, 6µM and 12µM) (lanes 2 to 4) for 12 h before cell lysis. Levels of CycT1 (1st panel), CDK9 (2nd panel) and the loading control actin (3rd panel) proteins were detected with anti-CycT1, anti-CDK9 and anti-β-actin antibodies, respectively, by WB. F. CycT1 levels in activated primary CD4+T cells are decreased by PKC inhibitors in a dose dependent manner. Activated primary CD4+T cells cells were untreated (lane 1) or treated with increasing amount of staurosporine (1.5µM, 3µM and 6µM) (lanes 2 to 4), bisindolylmaleinide IX (3µM, 6µM and 12µM) (lanes 5 to 7) for 12 h before cell lysis. Levels of CycT1 (1st panel), CDK9 (2nd panel) and the loading control actin (3rd panel) were detected with anti-CycT1, anti-CDK9 and anti-β-actin antibodies, respectively, by WB.

Additionally, co-IPs were performed in cells expressing the exogenous CycT1 or CycT1(280) proteins in the presence of bortezomib (12 h) and staurosporine (6 h) or bortezomib alone. As presented in Figs. 4A and 4B, interactions between exogenous CycT1 or CycT1(280) proteins and the endogenous CDK9 protein were significantly inhibited by staurosporine (1st panels, compare lanes 3 to lanes 2, ∼8.1-fold and ∼7.8-fold reduction, respectively). To confirm further that this PKC inhibitor could antagonize the phosphorylation of CycT1(280) by a high concentration of okadaic acid (Fig. 2C), cells expressing the exogenous CycT1(280) protein were treated with staurosporine for 6 h before adding okadaic acid for another 1.5 h, in the presence of bortezomib (12 h). Levels of threonine phosphorylation of CycT1(280) were compared by IPs with anti-HA antibodies. As presented in Fig. 4C, staurosporine antagonized completely the threonine phosphorylation of CycT1(280) (top two panels, compare lane 4 to lane 3).

Thus, PKC inhibitors, especially staurosporine, inhibit the phosphorylation of CycT1 and its interactions with CDK9, followed by the degradation of CycT1 in 293T cells.

To validate further the specificity of such negative regulation by PKC inhibitors, Jurkat and activated primary CD4+ T cells were also treated with different PKC inhibitors at increasing amounts for 12 h. As presented in Fig. 4D and compared to untreated cells, levels of endogenous CycT1 protein in Jurkat cells were significantly decreased by staurosporine in a dose dependent manner (1st panel, compare lanes 2, 3 and 4 to lane 1: 1.5µM, ∼2-fold; 3µM, ∼11-fold; 6µM, ∼20-fold reduction). Levels of CDK9 were largely unaffected (2nd panel in Fig. 4D). Similar to staurosporine, two additional PKC inhibitors, bisindolylmaleinide IX (Fig. 4E) and H-7 (Fig. S2B) also decreased levels of the endogenous CycT1 protein in a dose dependent manner. As presented in Fig. 4E, levels of CycT1 were decreased by bisindolylmaleinide IX ∼4-fold at 6µM and ∼10-fold at 12µM (1st panel, compare lanes 3 and 4 to lane 1). PKC inhibitor H-7 also decreased levels of CycT1 ∼6-fold at 70µM and ∼13-fold at 100µM (Fig. S2B, 1st panel, compare lanes 3 and 4 to lane 1). Levels of CDK9 were also largely unaffected by these inhibitors (Figs. 4D and S2B, 2nd panels).

Staurosporine and bisindolylmaleinide IX were also used in activated primary CD4+ T cells to confirm further our findings (Fig. 4F). In these cells, staurosporine depleted CycT1 up to 30-fold at 1.5, 3 and 6µM (donor 1, Fig. 4F, 1st panel, compare lanes 2, 3 and 4 to lane 1). Bisindolylmaleinide IX decreased them up to 20-fold at 3, 6 and 12µM (Fig. 4F, 1st panel, compare lanes 5, 6 and 7 to lane 1). In cells from donor 2, these compounds had similar effects on levels of CycT1 (Fig. S2C, 1st panel). Levels of CDK9 were largely unaffected by these inhibitors (Figs. 4F and S2C, 2nd panels). Activated primary CD4+ T cells were also treated with three additional PKC inhibitors (sotrastaurin, H-7, and HBDDE) for 12 h. As presented in Fig. S2D, all these PKC inhibitors decreased levels of CycT1 in a dose dependent manner (1st panel) without affecting those of CDK9 (2nd panel). Taken together, PKC inhibitors antagonize the phosphorylation of critical threonines in CycT1, which leads to the disassembly of P-TEFb and further degradation of CycT1.

### PKCα and PKCβ bind to CycT1, promote interactions between CycT1 and CDK9, and increase the stability of CycT1

Analysis of target specificities of our PKC inhibitors indicated that PKCα, PKCβ, PKCε represent candidate PKC isoforms responsible for the phosphorylation of CycT1. To validate that these PKC isoforms can target CycT1 for phosphorylation and promote P-TEFb assembly, their Flag-epitope-marked versions were expressed in 293T cells.

Different dominant (kinase) negative mutant PKC isoforms were also co-expressed with CycT1(280) or the mutant CycT1(280)TT143,149AA proteins in the presence of bortezomib for 12 h. Co-IPs were performed with anti-Flag antibodies. As presented in Fig. 5A, the mutant PKCαK368R (Soh & Weinstein, 2003) protein interacted with the mutant CycT1(280)TT143,149AA protein more potently than with CycT1(280) (1st panel, compare lanes 4 to 3, ∼3-fold increase), while no interactions with CDK9 were detected (Fig. 5A, 2nd panel, lanes 3 and 4). Since the catalytic domains of PKCβ1 and PKCβ2 are identical, and they differ only in their C-terminal 50 residues (Kubo, Ohno, & Suzuki, 1987). The dominant negative mutant PKCβ2K371R protein (Soh & Weinstein, 2003) was co-expressed with CycT1(280) or the mutant CycT1(280)TT143,149AA protein. Co-IPs were performed with anti-Flag antibodies. As presented in Fig. 5B, interactions between the mutant PKCβ2K371R protein and CycT1(280) or the mutant CycT1(280)TT143,149AA protein were detected (1st panel, compare lanes 4 to 3, ∼4.1-fold increase). Again, CDK9 did not interact with the mutant PKCβ2K371R protein (Fig. 5B, 2nd panel, lanes 3 and 4). In contrast, significantly reduced interactions were detected between CycT1(280) or the mutant CycT1(280)TT143,149AA proteins and PKCε, PKCδ, PKCγ and PKCθ (1st panels in Fig. S3A, S3B and S3C; data with PKCθ are not presented). We conclude that PKCα and PKCβ not only bind to but phosphorylate Thr143 and Thr149 in CycT1.

**Fig. 5.**
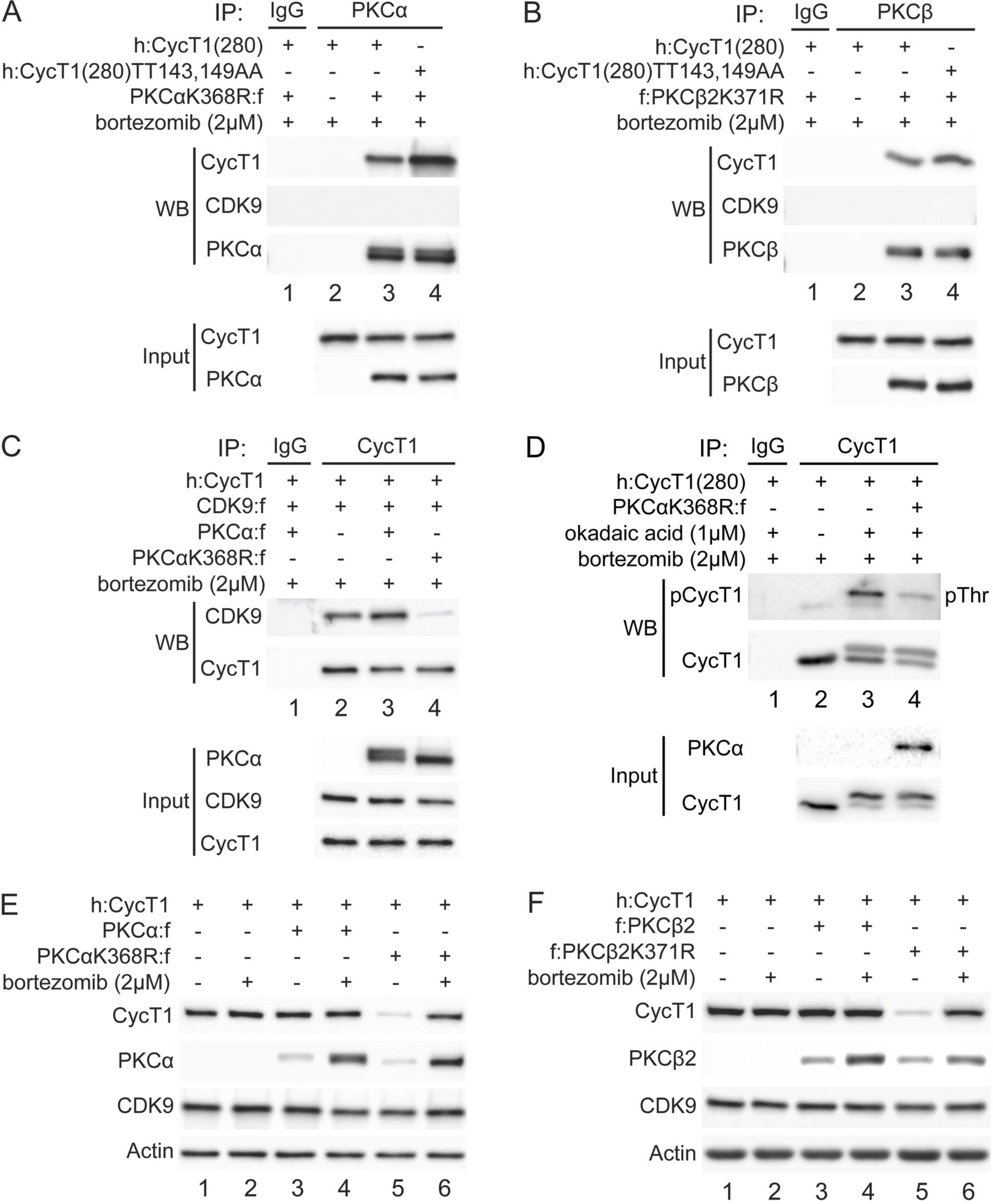
PKCα and PKCβ bind to CycT1 for its phosphorylation, also promote interactions between CycT1 and CDK9, and increase the stability of CycT1. A. PKCα binds to the truncated CycT1(280). Dominant negative mutant PKCαK368R protein and CycT1(280) or the mutant CycT1(280)TT143,149AA protein were co-expressed in the presence of 2µM bortezomib (+/− signs on top) in 293T cells. Co-IPs with PKCα are presented in the top three panels. 4th and 5th panels contain input levels of CycT1(280) and PKCα (input). B. PKCβ binds to CycT1(280). Dominant negative mutant PKCβ2K371R protein and CycT1(280) or the mutant CycT1(280)TT143,149AA protein were co-expressed in the presence of 2µM bortezomib (+/− signs on top) in 293T cells. Co-IPs with PKCβ are presented in the top three panels. 4th and 5th panels contain input levels of CycT1(280) and PKCβ (input). C. Dominant negative mutant PKCαK368R protein inhibits interactions between CDK9 and CycT1. PKCα or PKCαK368R, CycT1 and CDK9 were co-expressed in the presence of 2µM bortezomib (+/− signs on top) in 293T cells. Co-IPs with CycT1 are presented in top two panels. 3rd to 5th panels contain input levels of PKCα, CDK9 and CycT1 (input). D. PKCαK368R inhibits threonine phosphorylation of CycT1(280). CycT1(280) was expressed with or without PKCαK368R in the presence or absence of 1µM okadaic acid and 2µM bortezomib (+/− signs on top) in 293T cells. IPs with CycT1 were then probed with anti-pThr antibodies in the top panel. 3rd and 4th panels contain input levels of PKCα and CycT1(280) (input). E. PKCαK368R decreases levels of CycT1 in cells. PKCα or PKCαK368R, and CycT1 were co-expressed in the presence or absence of 2µM bortezomib (+/− signs on top) in 293T cells. Levels of CycT1 (1st panel), PKCα (2nd panel), CDK9 (3rd panel), and the loading control actin (4th panel) were detected with anti-HA, anti-Flag, anti-CDK9, and anti-β-actin antibodies, respectively, by WB. F. PKCβ2K371R decreases levels of CycT1. PKCβ2 or PKCβ2K371R, and CycT1 were co-expressed in the presence or absence of 2µM bortezomib (+/− signs on top) in 293T cells. Levels of CycT1 (1st panel), PKCβ2 (2nd panel), CDK9 (3rd panel), and the loading control actin (4th panel) were detected with anti-HA, anti-Flag, anti-CDK9, and anti-β-actin antibodies, respectively, by WB.

To confirm further that PKCα and PKCβ contribute to the assembly and stability of P-TEFb, PKCα or the mutant PKCαK386R protein were co-expressed with CycT1(280) and CDK9 in the presence of bortezomib (12 h) in 293 T cells. Interactions between these proteins were analyzed by co-IPs with anti-HA antibodies. As presented in Fig. 5C, while PKCα slightly increased CycT1:CDK9 interactions (Fig. 5C, 1st panel, compare lane 3 to lane 2), the mutant PKCαK386R protein inhibited them (1st panel, compare lane 4 to lanes 2 and 3, ∼7.9-fold reduction). To confirm that decreased interactions between CycT1(280) and CDK9 by the mutant PKCαK386R protein were caused by the inhibition of PKC-dependent phosphorylation of CycT1(280), it was co-expressed with the mutant PKCαK386R protein in the presence of bortezomib (12 h) and okadaic acid (1.5 h). IPs were conducted with anti-HA antibodies. As presented in Fig. 5D, the expression of the mutant PKCαK386R protein decreased levels of threonine phosphorylation in CycT1(280) by ∼5.2-fold as detected with anti-pThr antibodies (1st panel, compare lanes 4 to 3). To demonstrate if the mutant PKCαK386R protein also decreased levels of CycT1 protein, PKCα or the mutant PKCαK386R protein were co-expressed with the CycT1 protein in the presence or absence of bortezomib (12 h). As presented in Fig. 5E, CycT1 co-expressed with PKCα had similar levels of expression as CycT1 without PKCα co-expression, which was not affected by bortezomib (1st panel, compare lanes 3 to 1; compare lanes 3 to 4; compare lanes 1 to 2). In sharp contract, co-expression of the mutant PKCαK386R protein decreased greatly levels of CycT1 protein in these cells (Fig. 5E, 1st panel, compare lane 5 to lanes 1 and 3, ∼9.1-fold reduction), which was reversed by bortezomib (Fig. 5E, 1st panel, compare lane 6 to lane 5). Similar to PKCαK386R, co-expressed PKCβ2K371R also significantly diminished levels of CycT1 protein (Fig. 5F, 1st panel, compare lane 5 to lanes 1 and 3, ∼10.4-fold reduction), which was reversed by bortezomib (Fig. 5F, 1st panel, compare lane 6 to lane 5). Levels of PKCα, PKCαK386R, PKCβ2, and PKCβ2K371R were increased by bortezomib, which is consistent with the demonstrated instability of PKC (Lu et al., 1998) (Fig. 5E and 5F, 2nd panels, compare lanes 3 and 5 to lanes 4 and 6). Also, levels of the endogenous CDK9 protein were unaffected in these cells (Fig. 5E and 5F, 3rd panels). Taken together, CycT1 is targeted mainly by PKCα and PKCβ, which not only promote interactions between CycT1 and CDK9 via the phosphorylation of CycT1, but also stabilize CycT1 in cells.

### Depletion of PKC leads to decreased levels of CycT1 in cells

Previous papers demonstrated that isoforms of PKC are inactive or absent in resting cells (Heissmeyer et al., 2004; Pfeifhofer-Obermair, Thuille, & Baier, 2012). Moreover, phorbol esters (PMA) deplete PKC in most cells (Manger, Weiss, Imboden, Laing, & Stobo, 1987). In addition, HIV Tat, whose proteome first identified and whose co-activator is P-TEFb, no longer works in these cells (Jakobovits, Rosenthal, & Capon, 1990). Together with our data, it appears that PKC influences dynamic changes of P-TEFb in different cell types and under varying conditions. To examine this situation further, 100ng/ml PMA was administered to Jurkat cells or activated primary CD4+ T cells for several days. As presented in Fig. 6A, endogenous CycT1 protein levels were decreased up to ∼6-fold at 72 h and ∼11-fold at 96 h after the addition of PMA (1st panel, compare lanes 2 and 3 to lane 1). Levels of CDK9 protein were largely unaffected (Fig.6A, 2nd panel, lanes 1 to 3). Furthermore, the same PMA treatment was performed in activated primary CD4+ T cells from two different donors. Similar to Jurkat cells, activated primary CD4+ T cells from donor 1 lost CycT1 expression up to ∼7-fold at 72 h and ∼16-fold at 96 h after PMA treatment (Fig. 6B, 1st panel, compare lanes 2 and 3 to lane 1). Again, levels of CDK9 were largely unaffected (Fig.6B, 2nd panel, lanes 1 to 3). Levels of PKCα were equivalently decreased at 72 and 96 h after the addition of PMA (Fig. 6B, 3rd panel, compare lanes 2 and 3 to lane 2). Other PKC isoforms, PKCβ1 and PKCβ2 were also depleted at these time points (Fig. 6B, 4th and 5th panels, compare lanes 2 and 3 to lane 2). These cells do not express PKCε (Fig. 6B, 6th panel, lanes 1 to 3). Same changes were also observed in activated primary CD4+ T cells from donor 2 (Fig. S4A). We also found that the addition of bortezomib for another 24 h after 72 h PMA incubation rescued most of these decreased levels of CycT1 in activated primary CD4+ T cells (Fig. 6C, 3rd panel, compare lane 3 to lane 2). Co-IPs were also conducted in cells treated with bortezomib alone or PMA and bortezomib with anti-CDK9 antibodies. As presented in Fig. 6C, interactions between CycT1 and CDK9 were significantly decreased in PMA-treated cells (up to ∼7.6-fold), compared to controls (Fig. 6C, 1st panel, compare lane 3 to lane 2). These data demonstrate that the depletion of PKC in Jurkat cells and activated primary CD4+ T cells by PMA treatment causes the dissociation of P-TEFb and depletion of CycT1. This finding explains the hereto puzzling observation that Tat does not work in cells treated with PMA (Jakobovits et al., 1990).

**Fig. 6.**
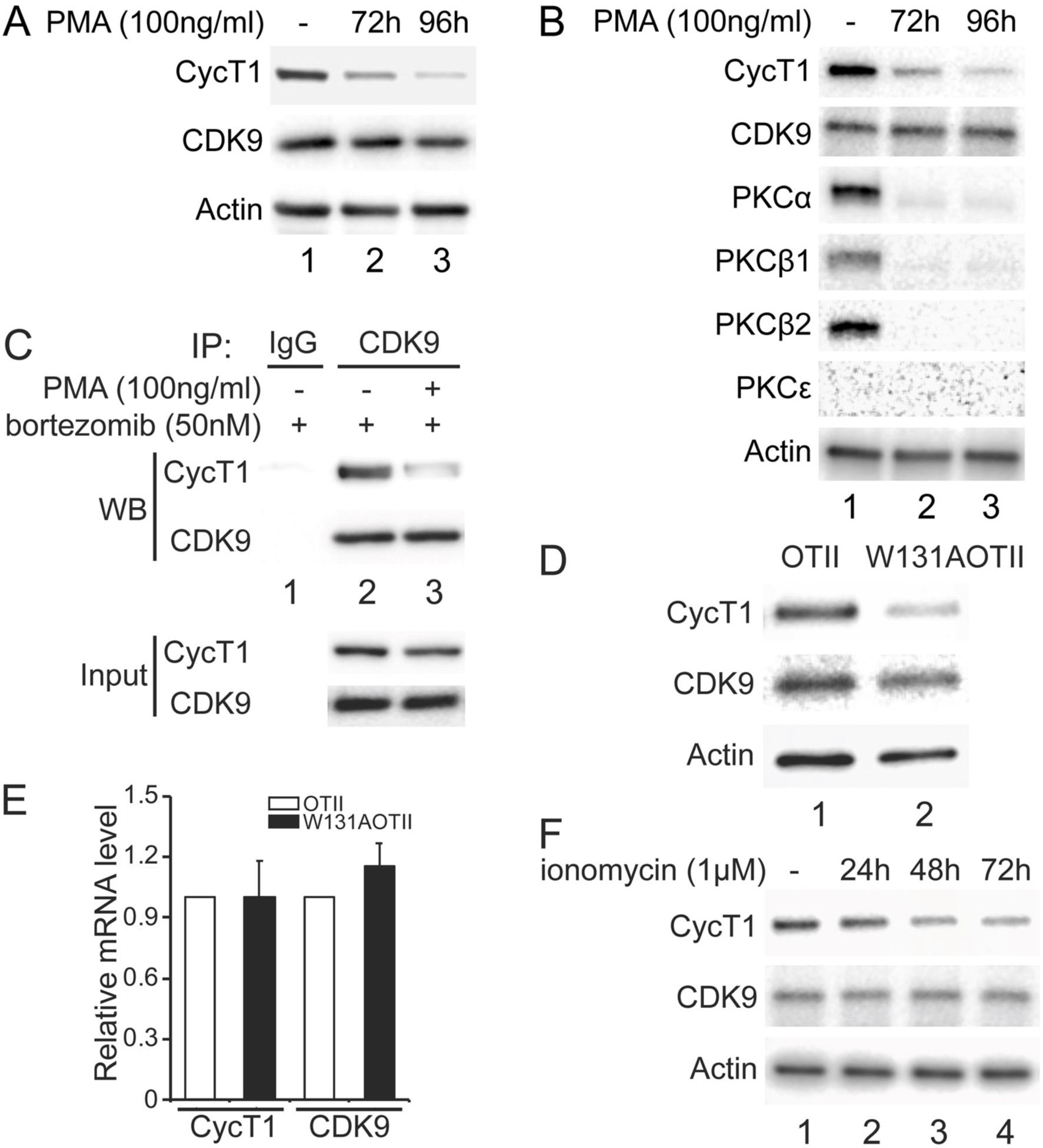
Depletion of PKCs leads to decreased levels of CycT1 in cell lines and primary cells. A. Prolonged PMA treatment decreases levels of CycT1 in Jurkat cells. Jurkat cells were untreated (lane 1) or treated with 100 ng/ml PMA for 72 and 96 h (lane 2 and 3) before cell lysis. Top panels contain levels of endogenous CycT1 and CDK9 proteins, the bottom panel contains the loading control actin protein. B. Prolonged PMA treatment decreases levels of CycT1 and PKC in activated primary CD4+ T cells. Activated primary CD4+ T cells were untreated (lane 1) or treated with 100 ng/ml PMA for 72 and 96 h (lanes 2 and 3) before cell lysis. Upper panels contain levels of endogenous CycT1, CDK9, PKCα, PKCβ1, PKCβ2, and PKCε proteins. The bottom panel contains the loading control actin protein. C. Depletion of PKC impairs interactions between CycT1 and CDK9 in activated primary CD4+ T cells. Activated primary CD4+ T cells were treated with or without 100 ng/ml PMA (+/− signs on top) for 96 h. At 72 h, 50nM bortezomib was added for additional 24 h before cell lysis. Co-IPs with CDK9 are presented in top two panels. Panels 3 and 4 contain input CycT1 and CDK9 proteins (input). D. CycT1 levels are decreased in mouse anergic T cells. T cells were selected from WT OTII (WT ZAP70) or mutant W131AOTII (ZAP70W131A) mice and lysed. Upper panels contain levels of endogenous CycT1 and CDK9 proteins. Lanes are: lane 1, WT OTII mice; lane 2, mutant W131AOTII mice. Bottom panel contains the loading control actin protein. E. mRNA levels of CycT1 and CDK9 are equal in mouse anergic and WT T cells. Relative mRNA levels of CycT1 (Left 2 bar graphs) and CDK9 (right 2 bar graphs) are presented as -fold change in W131AOTII T cells (black bars) above levels of WT OTII T cells (white bars). Error bars represent S.E., n = 3. F. Prolonged ionomycin treatment decreases levels of CycT1 in activated primary CD4+ T cells. Activated primary CD4+ T cells were untreated (lane 1) or treated with 1µM ionomycin for 24, 48 and 72 h (lanes 2 to 4) before cell lysis. Upper 2 panels contain levels of endogenous CycT1 and CDK9 proteins. The bottom panel contains the loading control actin protein.

We observed previously that levels of CycT1 increase significantly in resting CD4+ T cells with the addition of bortezomib (Cary & Peterlin, 2020). Nevertheless, interactions between CycT1 and CDK9 remain lower than in activated primary CD4+ T cells. To extend these findings to anergic T cells that lose the ability to respond to agonist antigen or stimulation of the T cell antigen receptor, we examined W131AOTII T cells from mice where the endogenous ZAP70 protein was substituted by a constitutively active mutant ZAP70-W131A protein. Introduction of the W131A mutant into the OTII transgenic background (W131AOTII) results in high numbers of anergic and CD4 regulatory T cells (Palacios & Weiss, 2007). As presented in Fig. 6D, levels of CycT1 protein were significantly lower in W131AOTII than in control OTII T cells (1st panel, compare lane 2 to lane 1, ∼7.8-fold decrease). Levels of the CDK9 were largely unchanged in these cells (Fig. 6D, 2nd panel, compare lane 2 to lane 1). Meanwhile, levels of CycT1 and CDK9 transcripts in W131AOTII and OTII cells remained unchanged (Fig. 6E), which is consistent with previous observations that mRNA levels of CycT1 and CDK9 do not vary between resting and activated CD4+ T cells (Cary & Peterlin, 2020; Sung & Rice, 2006). Moreover, since these W131AOTII T cells exhibit impaired T cell receptor signaling, activating these cells with anti-CD3 and anti-CD28 antibodies did not increase levels of CycT1 (data not presented). It was demonstrated that treatment of T cells with calcium ionophores such as ionomycin also depletes PKC and induces anergy (Heissmeyer et al., 2004). Therefore, we examined whether sustained ionomycin treatment of primary activated CD4+ T cells causes depletion of CycT1. As presented in Fig. 6F, CycT1 expression in activated primary T cells from a donor began to decrease at 24 h (1st panel, compare lanes 2 to 1, ∼1.8-fold reduction) after the addition of ionomycin (1µM), and continued at 48 h (1st panel, compare lanes 3 to 1, ∼4.5-fold reduction) and 72 h (1st panel, compare lanes 4 to 1, ∼9.1-fold reduction). Levels of CDK9 were largely unaffected (Fig. 6F, 2nd panel, lanes 1 to 3). Similar changes were also observed with donor 2 treated with ionomycin (Fig. S4B). Taken together, the absence of active PKC correlates with significant decreases of CycT1, which prevents the assembly of the functional P-TEFb complex.

## Discussion

In this study, we found that mutations of two critical residues in CycT1 (Thr143 and Thr149 to alanine) impair its binding to CDK9. Phosphorylation of these residues is required to promote CycT1:CDK9 interactions. Structural analyses revealed that phosphates on Thr143 and Thr149 in CycT1 increase intramolecular and intermolecular binding to specific residues in CycT1 and CDK9, respectively, which potentiates P-TEFb assembly and stabilizes CycT1. This prediction was confirmed experimentally. Thr143 and Thr149 are located in PKC consensus sequences. Indeed, PKC inhibitors inhibit CycT1 phosphorylation. PP1 then dephosphorylates CycT1. As a consequence, interactions between CycT1 and CDK9 are attenuated, CycT1 is rapidly degraded and P-TEFb disappears. Of PKC isoforms, PKCα and PKCβ not only bind to CycT1 but promote CycT1:CDK9 interactions and stabilize CycT1. Finally, depleting PKC or its inactivity leads to CycT1 dissociation from CDK9 and its degradation in transformed cell lines as well as primary activated and anergic T cells. We conclude that the assembly and stability of P-TEFb require the phosphorylation of CycT1, which is regulated by PKCα/PKCβ and PP1.

Ours is the first study that addresses the reversible phosphorylation of CycT1. Based on previous mutageneses of CycT1, three threonine residues (Thr143, 149 and 155) could affect CycT1 stability and P-TEFb assembly. They are conserved in over 142 mammalian species and in CycT2 (sequences are picked out from NCBI and aligned).

Moreover, previous reports indicated that these residues play critical roles in P-TEFb function (Jadlowsky, Nojima, Okamoto, & Fujinaga, 2008; Kuzmina et al., 2014). Of these, we identified Thr143 and Thr149 to be responsible for P-TEFb assembly and CycT1 stability. Their dephosphorylation was blocked by high dose okadaic acid, which inhibits PP1. PP1 contains three catalytic and over fifty regulatory subunits, whose genetic inactivation is lethal to cells (Cohen, 2002; Ferreira, Beullens, Bollen, & Van Eynde, 2019). Thus, phosphatase inhibitor okadaic acid (low dose PP2A, high dose PP1) is used to distinguish between PP1 or PP2. In other studies, they defined phosphatases that regulate cell cycle cyclins, such as CycB, CycD1 and AIB1 (Edelson & Brautigan, 2011; Ferrero et al., 2011; Vorlaufer & Peters, 1998). As to kinases, PKCα and PKCβ isoforms phosphorylate these two threonine residues in CycT1 for P-TEFb assembly. This finding was confirmed by PKC inhibitors, direct binding studies and the use of dominant kinase negative mutant PKC proteins. Since its degradation and assembly have to occur in most if not all cells of the organism, the involvement of ubiquitously expressed and reductant kinases and phosphatases in this post-translational regulation of P-TEFb is not unexpected. Importantly, our study also reveals that the depletion of PKC, as occurs after chronic activation or phorbol ester treatment, results in P-TEFb disassembly and CycT1 degradation, which explains cells becoming unresponsive to external stimuli. It also explains why Tat, whose co-activator is P-TEFb, no longer functions in such cells (Jakobovits et al., 1990). Anergic cells also lack P-TEFb (Fig. 6D), which contributes to their unresponsive phenotype. From our and other studies, we also know that resting and memory T cells lack P-TEFb (Garriga et al., 1998; Ghose et al., 2001). Thus, proviral latency is maintained in these cells (Fujinaga & Cary, 2020). This situation most likely pertains to other DNA viruses, such as herpes simplex virus and HTLV1/2 in yet other resting or terminally differentiated cells (Kulkarni & Bangham, 2018; Nicoll, Proenca, & Efstathiou, 2012).

Elegant studies revealed that T-loop phosphorylation (Thr186) and of Ser175 in CDK9, are important for the activity of P-TEFb (Zhou et al., 2012). Global analyses also revealed two phosphorylation sites in the C-terminal half of CycT1 (Mbonye et al., 2013). However Thr143 and Thr149 were missed, possibly due to the metabolic state of examined cells or lack of adequate phosphatase inhibition. Unphosphorylated CycT1 dissociates from P-TEFb and is degraded. CDK9 remains and is stabilized by HSP70 and HSP90 (O’Keeffe et al., 2000). This observation presents uncanny similarities with the regulation of cell cycle CDKs, whose levels are also regulated by reversible phosphorylation, disassembly and degradation. For example, the phosphorylation of CycE determines its stability. Only upon its dephosphorylation, dissociation and degradation can the cell cycle proceed (Clurman, Sheaff, Thress, Groudine, & Roberts, 1996). Thus, the transcriptional CDKs display and mimic regulatory paradigms of cell cycle CDKs. This regulation is also very different from the P-TEFb equilibrium in growing, proliferating cells, where 7SK snRNP plays the major role. There, P-TEFb partitions between the active free state, bound to activators and/or the super-elongation complex, and the inactive state, where 7SK snRNA coordinates its sequestration with HEXIM1/2, LaRP7, MePCE in the 7SK snRNP. In this RNP, HEXIM1 or HEXIM2 inhibit CDK9 by binding to its ATP pocket, a situation that is reminiscent of CDK2 inhibition by p21 (Russo, Jeffrey, Patten, Massague, & Pavletich, 1996). Thus, our study has revealed an additional important aspect of the regulation of transcriptional elongation in cells.

Global mRNA sequencing studies examine changes in levels of specific transcripts between and in different states, i.e. activation, proliferation and differentiation of cells (Sung & Rice, 2009). However, levels of CycT1 and CDK9 mRNAs do not vary between resting and activated T cells (Marshall, Salerno, Garriga, & Grana, 2005), between responsive and anergic T cells or exhausted T cells. This finding is important, as these global studies do not look at levels of resulting proteins. Thus, they underestimate contributions of P-TEFb and/or other transcriptional CDKs in their evaluations of target genes. Additionally, P-TEFb must be recruited by transcription factors to the paused RNAPII at promoters. Indeed, P-TEFb can mediate short and long long-distance interactions between enhancers and promoters to promote transcription elongation and co-transcriptional processing of individual and clusters of genes (Taube et al., 2002). In this scenario, the C-terminal His-rich region of CycT1 interacts directly with the CTD of RNAPII (Taube et al., 2002). Thus, these post-translational modifications of P-TEFb play critical roles in activated transcription and for growth and proliferation of cells. In quiescent cells, only basal transcription is detected (Cheung & Rando, 2013; Roche, Arcangioli, & Martienssen, 2017). Importantly, external stimuli can reverse this phenotype via this reversible phosphorylation of P-TEFb. As a result, effects of potent activators, such as cMyc, NF-kB, steroid hormones, CIITA, etc are translated to the productive transcription of their target genes (Fujinaga, 2020).

Finally, since our study revealed families of kinases and phosphatases that affect P-TEFb, it is possible that the use of more targeted phosphatase inhibitors could block this transition to quiescence and terminal differentiation of cells. Their use might even prevent the establishment of proviral latency. Although the substitution of phosphomimetic residues for serine/threonine/tyrosine residues is not always successful, it is possible that such modifications in CycT1 could create a constitutively active P-TEFb complex. Additionally, target residues on CycT1 and CDK9 could be changed to stabilize this structure. If successful, modified P-TEFb complexes could be studied for effects on immune responses as well as latency induction and reversal in many different scenarios. The other approach would be to identify the E3 ligase that is responsible for the degradation of the unphosphorylated CycT1 protein. To this end, bortezomib and other proteasomal inhibitors have been examined already for the reversal of HIV latency in resting CD4+ T cells (Cary & Peterlin, 2020; Li et al., 2019).

Taking all these findings into account, the understanding of complex post-translational regulation of P-TEFb promises to reveal additional approaches not only to proviral latency, but to host immune responses, cellular regeneration and dedifferentiation.

## Materials and Methods

### Plasmids, Reagents and Antibodies

HA-CycT1 (h:CycT1), CDK9-Flag (CDK9:f) and plasmids containing mutated CycT1 or CDK9 sequences were constructed by cloning PCR fragments containing the coding sequences of CycT1 and CDK9 into pcDNA3.1 vector with indicated epitope tags. PKC plasmids (PKCα, β, γ, δ, ε and θ) were obtained from Addgene, and their coding sequences were subcloned into the pcDNA3.1 vector containing the Flag epitope tag. All the reagents and antibodies are listed in the table within the supplementary data.

### Cell culture

Human Embryonic Kidney (HEK) 293T cells were cultured in Dulbecco’s modified Eagle’s medium (DMEM) (Corning) with 10% fetal bovine serum (FBS) (Sigma Aldrich), Jurkat cells and Peripheral Blood Mononuclear Cells (PBMCs) were cultured in Roswell Park Memorial Institute (RPMI) 1640 (Corning) with 10% FBS at 37 °C and 5% CO2.

Resting CD4+ T cells were purified from bulk PBMCs by using Dynabeads™ Untouched™ Human CD4 T Cells Kit (ThermoFisher Scientific). Selected CD4+ T cells were activated by Dynabeads™ Human T-Activator CD3/CD28 kit (ThermoFisher Scientific) and were maintained in RPMI 1640 with 10% FBS, containing 30U/ml IL-2.

### Cell manipulation

Transfection of plasmid DNA was conducted in 293T cells using Lipofectamine 3000 (Life Technology) and X-tremeGENE™ HP DNA Transfection Reagent (Roche) according to the manufacturer’s instructions.

293T cells were treated with 2.5µM bortezomib for 12 h, Jurkat cells were treated with 100nM bortezomib for 12 h, and activated CD4+ T cells were treated with 50nM bortezomib for 12 h before the cell lysis. 293T cells were treated with 5nM or 1µM okadaic Acid for 1.5 h before the cell lysis. 293T cells, Jurkat cells and activated CD4+ T cells were treated with different PKC inhibitors (sotrastaurin, H-7, staurosporine, Go 6976 and bisindolylmaleinide IX) for 6 h or 12 h before the cell lysis.

### Co-immunoprecipitation (Co-IP) and protein Densitometry

293T, Jurkat or CD4+ T cells were lysed on ice using RIPA buffer (50mM Tris-HCl, pH 8.0, 5 mM EDTA, 0.1% SDS, 1.0% Nonidet P-40, 0.5% sodium deoxycholate, 150 mM NaCl) supplemented with the protease and phosphatase inhibitors, then for one time sonication (level 4, 2 s), followed by a 10 min centrifugation (21,000×g). The supernatant was precleared, and incubated with indicated primary antibodies or control IgG overnight. Mixtures were then incubated with protein G-Sepharose beads for additional 2h, followed by 5 times’ wash with RIPA buffer (500mM NaCl). Co-IP samples and input (1% of whole cell lysates) were subjected to western blotting (WB) as described previously (Huang, Shao, Fujinaga, & Peterlin, 2018).

Protein densitometry was obtained using Image Studio software (LI-COR). Relative protein expression in whole cell lysates was calculated by normalizing the indicated proteins with loading control β-actin. Quantification data were presented as fold change over values obtained with control samples.

### W131AOTII and control OTII cells preparation

W131AOTII mice were described previously (Hsu, Cheng, Chen, Liang, & Weiss, 2017). Control OTII TCR transgenic mice were purchased from The Jackson Laboratory.

Peripheral naïve (CD44^low^CD62L^+^) CD25^-^Va2^+^CD4^+^ T cells were sorted from combined lymphoid organs (spleens and lymph nodes) of OTII or W131AOTII mice (8–12 weeks of age). The cells (10^6^ cells) were washed with PBS, the supernatant was aspirated, and the pelleted cells were lysed as described above, then subjecting into WB assay.

## Acknowledgement

This study was supported by: Damon Runyon Cancer Research Foundation Fellowship (to T.T.T.N.); NIH R01 AI049104 (F.H., D.C.C., H.P., B.M.P., and K.F.); NIH P01 AI091580 (to T.T.T.N. and A.W.); Howard Hughes Medical Institute (to T.T.T.N. and A.W.); Nora Eccles Treadwell Foundation. (F.H., H.P., and K.F.); HARC center (NIH P50AI150476) (to F.H., D.C.C., H.P., B.M.P., K.F., R.R., I.E., and A.S.). We thank Zeping Luo (lab member) for excellent technical assistance.

## Author contribution

F.H. designed and conducted experiments, interpreted the data, and wrote the manuscript. K.F and B.M.P supervised experiments, helped to interpret results and to write the manuscript. R.R., I.A., and A.S. performed structural analyses. T.T.T.N. prepared OTII and W131AOTII lysates and performed RNAsequencing. T.T.T.N. and A.W. also interpreted data from anergic T cells. H.P. and D.C.C. selected primary CD4+ T cells. All authors contributed to the writing and editing of the manuscript.

## Competing Interest Statement

The authors declare no competing financial interests.

## Supplementary data

Reagents and Antibodies

| REAGENT | MANUFACTURER | IDENTIFIER |
| --- | --- | --- |
| **ANTIBODIES** |  |  |
| mouse monoclonal anti-CycT1 | Santa Cruz Biotechnology | E-3, sc-271348 |
| mouse monoclonal anti-CDK9 | Santa Cruz Biotechnology | F-6, sc-376646 |
| mouse monoclonal anti-PKCα | Santa Cruz Biotechnology | H-7, sc-8393 |
| mouse monoclonal anti-PKCβ1 | Santa Cruz Biotechnology | E-3, sc-8049 |
| mouse monoclonal anti-PKCβ2 | Santa Cruz Biotechnology | F-7, sc-13149 |
| mouse monoclonal anti-PKCε | Santa Cruz Biotechnology | E-5, sc-1681 |
| rabbit polyclonal anti-CDK9 | Abcam | EPR22956-37, ab239364 |
| mouse monoclonal anti-HA | Sigma-Aldrich | H3663 |
| mouse monoclonal anti-Flag | Sigma-Aldrich | F1804 |
| rabbit monoclonal anti-HA | Sigma-Aldrich | H6908 |
| rabbit monoclonal anti-Flag | Sigma-Aldrich | F7425 |
| rabbit monoclonal anti-CycT1 | Cell Signaling Technology | D1B6G, #81464S |
| rabbit monoclonal anti-β-Actin | Cell Signaling Technology | 13E5, #4970S |
| mouse monoclonal anti-Phospho-Threonine | Cell Signaling Technology | 42H4, #9386S |
| normal rabbit control IgG | Santa Cruz Biotechnology | sc-2027 |
| normal mouse control IgG | Santa Cruz Biotechnology | sc-2050 |
| ECL mouse IgG HRP-linked whole Ab | GE Healthcare | NA9310 |
| ECL rabbit IgG HRP-linked whole Ab | GE Healthcare | NA9340 |
| **CHEMICALS** |  |  |
| MG132 | Sigma-Aldrich | M7449 |
| bortezomib | Calbiochem | 179324-69-7 |
| okadaic acid | Cell Signaling Technology | #5934S |
| staurosporine | Selleckchem | S1421 |
| sotrastaurin | Selleckchem | S2791 |
| Go 6976 | Selleckchem | S7119 |
| bisindolylmaleinide IX | Selleckchem | S7207 |
| HBDDE | Selleckchem | ab141573 |
| H-7 | Abcam | ab142308 |
| cycloheximide (CHX) | Sigma-Aldrich | C4859 |
| interleukin-2 (IL-2) | Sigma-Aldrich | I7908 |
| phorbol 12-myristate 13-acetate (PMA) | Sigma-Aldrich | P8139 |
| ionomycin | Sigma-Aldrich | I9657 |

**Fig. S1.**
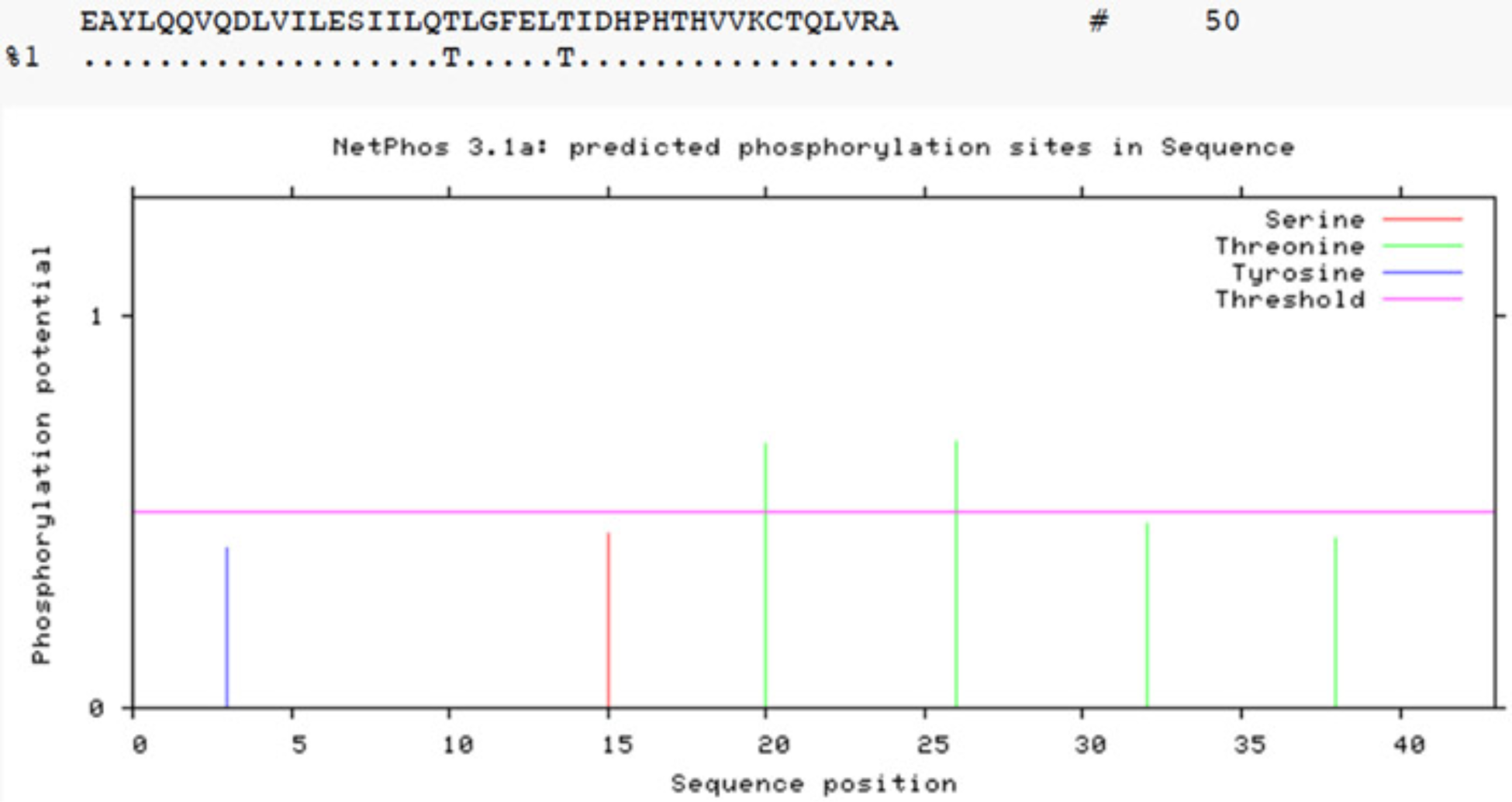
Thr143 and T149 are the phosphorylated residues in CycT1. Potential phosphorylation sites between 124 aa. to 166 aa. In CycT1 were predicted by the NetPhos 3.1 program. Threshold was set to 0.5 (default value), indicated by the pink line. Thr143 and Thr149 scored highest for potential phosphorylation sites.

**Fig. S2.**
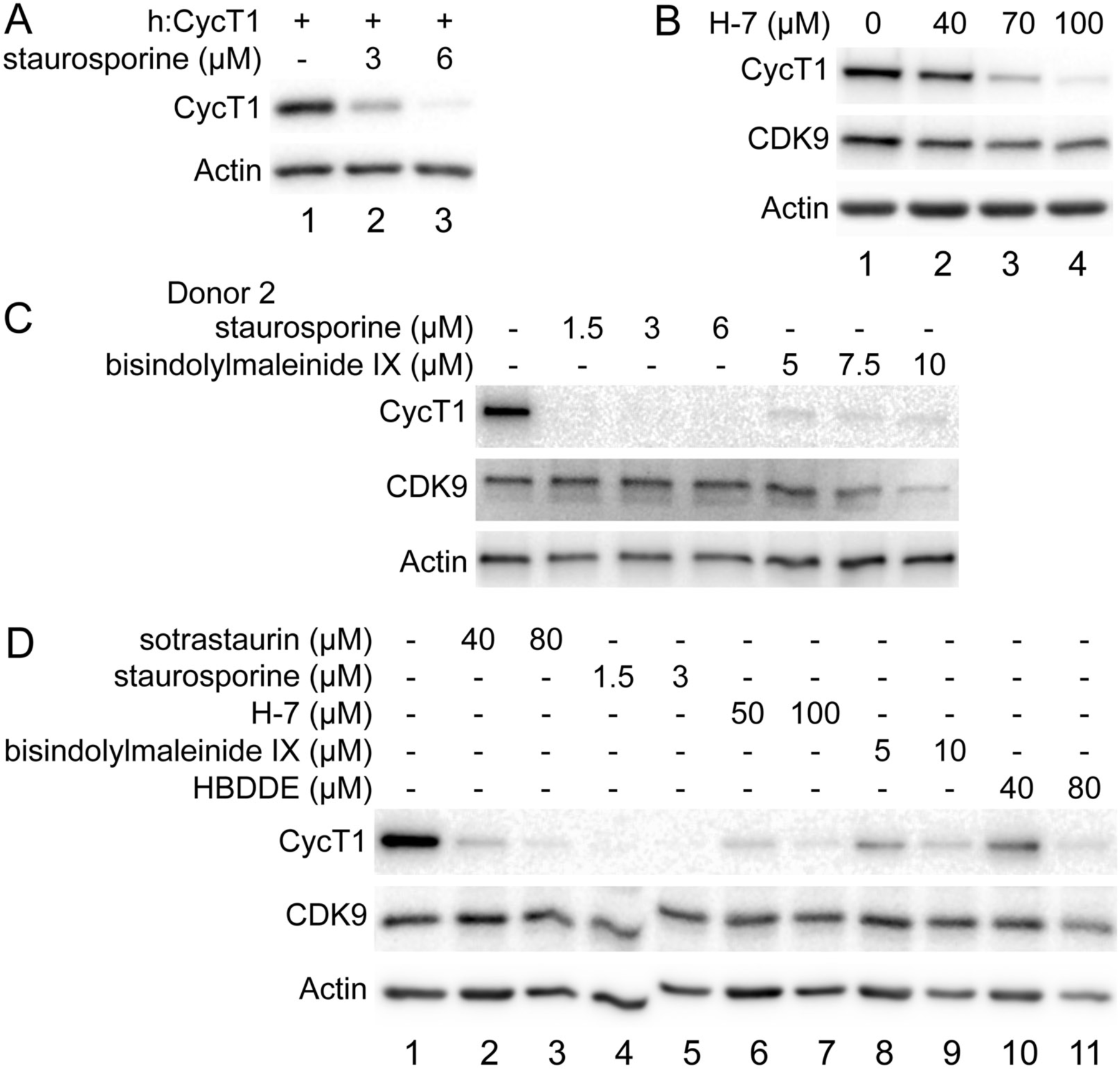
PKC inhibitors promote CycT1 degradation in different cells. A. Staurosporine decreases exogenous CycT1 levels in a dose dependent manner. 293T cells expressing CycT1 were untreated (lane 1) or treated with increasing doses of staurosporine (3µM and 6µM) (lanes 2 and 3) for 12 h before cell lysis. Levels of CycT1 (1st panel) and the loading control actin (2nd panel) protein were detected with anti-HA and anti-β-actin antibodies, respectively, by WB. B. PKC inhibitor H-7 decreases CycT1 levels in a dose dependent manner. Jurkat cells were untreated (lane 1) or treated with increasing doses of bisindolylmaleinide IX (40µM, 70µM and 100µM) (lanes 2 to 4) for 12 h before cell lysis. Levels of CycT1 (1st panel), CDK9 (2nd panel) and the loading control actin (3rd panel) protein were detected with anti-CycT1, anti-CDK9 and anti-β-actin antibodies, respectively, by WB. C. CycT1 levels in activated primary CD4+T cells (donor 2) are decreased by PKC inhibitors in a dose dependent manner. Activated primary CD4+T cells cells were untreated (lane 1) or treated with increasing amount of staurosporine (1.5µM, 3µM and 6µM) (lanes 2 to 4), bisindolylmaleinide IX (3µM, 6µM and 12µM) (lanes 5 to 7) for 12 h before cell lysis. Levels of CycT1 (1st panel), CDK9 (2nd panel) and the loading control actin (3rd panel) protein were detected with anti-CycT1, anti-CDK9 and anti-β-actin antibodies, respectively, by WB. D. Five different PKC inhibitors decrease levels of CycT1 in activated primary CD4+T cells in a dose dependent manner. Activated primary CD4+T cells cells were untreated (lane 1) or treated with increasing amounts of sotrastaurin (40µM and 80µM) (lanes 2 and 3), staurosporine (1.5µM and 3µM) (lanes 4 and 5), H-7 (50µM and 100µM) (lanes 6 and 7), bisindolylmaleinide IX (5µM and 10µM) (lane 8 and lane 9) and HBDDE (40µM and 80µM) (lanes 10 and 11) for 12 h before cell lysis. Levels of CycT1 (1st panel), CDK9 (2nd panel) and the loading control actin (3rd panel) protein were detected with anti-CycT1, anti-CDK9 and anti-β-actin antibodies, respectively, by WB.

**Fig. S3.**
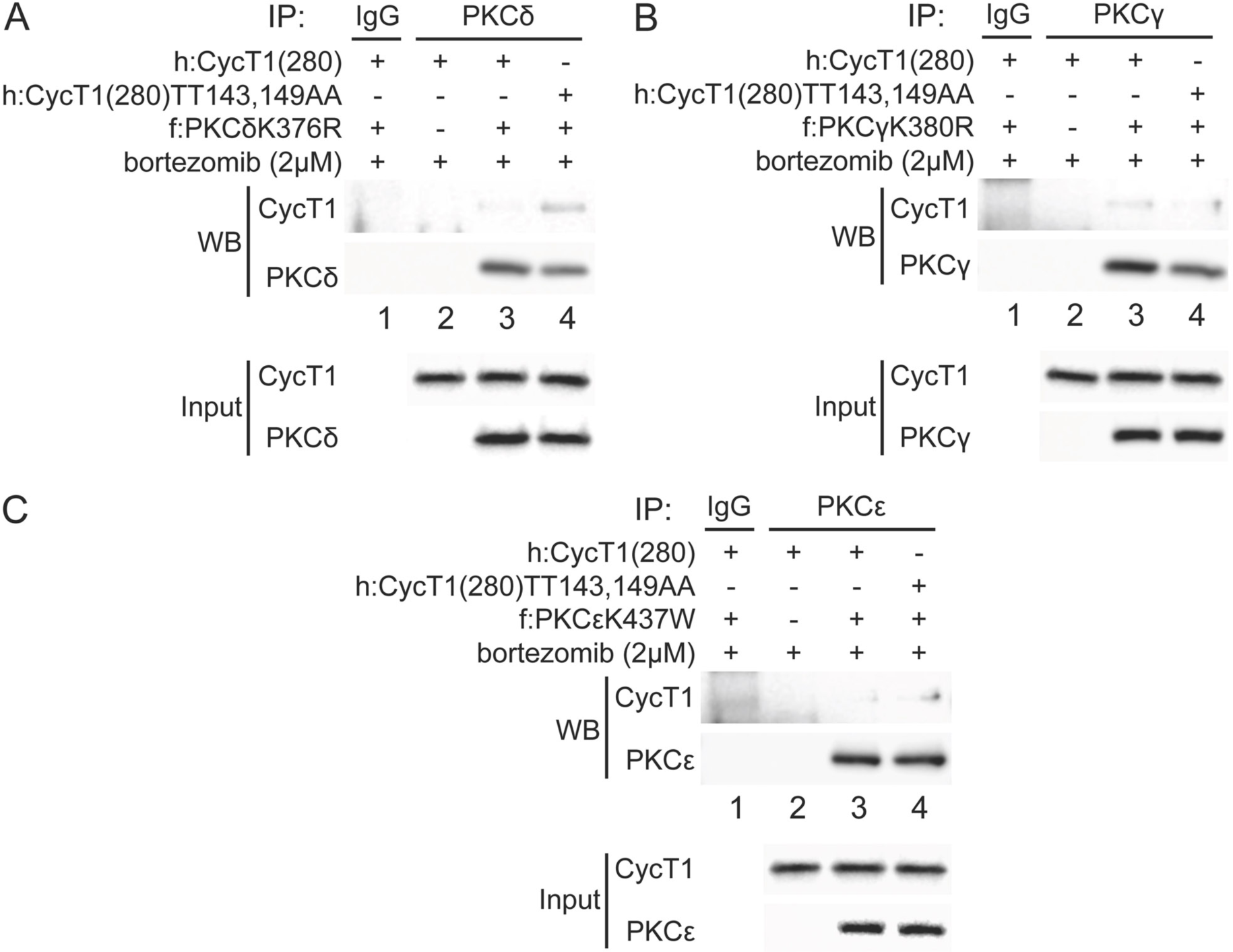
PKCδ, PKCγ, and PKCε bind weakly to CycT1. A. PKCδ binds weakly to CycT1(280). Dominant negative mutant PKCδK376R proteins and CycT1(280) or the mutant CycT1(280)TT143,149AA protein were co-expressed in the presence of 2µM bortezomib in 293T cells. Co-IPs with PKCδ are presented in the top two panels. Lanes are: lane 1, IgG control; lane 2, CycT1(280); lane 3 CycT1(280) and the mutant PKCδK376R protein; lane 4, mutant CycT1(280)TT143,149AA and PKCδK376R proteins. 3th and 4th panels contain input levels of CycT1(280) and PKCδ (input). B. PKCγ binds weakly to CycT1(280). Dominant negative mutant PKCγK380R protein and CycT1(280) or the mutant CycT1(280)TT143,149AA protein were co-expressed in the presence of 2µM bortezomib in 293T cells. Co-IPs with PKCγ are presented in the top two panels. Lanes are: lane 1, IgG control; lane 2, CycT1(280); lane 3 CycT1(280) and the mutant PKCγK380R protein; lane 4, mutant CycT1(280)TT143,149AA and PKCγK380R proteins. 3th and 4th panels contain input levels of CycT1(280) and PKCγ (input). C. PKCε binds weakly to CycT1(280). Dominant negative mutant PKCεK437W protein and CycT1(280) or the mutant CycT1(280)TT143,149AA protein were co-expressed in the presence of 2µM bortezomib in 293T cells. Co-IPs with PKCε are presented in the top two panels. Lanes are: lane 1, IgG control; lane 2, CycT1(280); lane 3 CycT1(280) and the mutant PKCεK437W protein; lane 4, mutant CycT1(280)TT143,149AA and PKCεK437W proteins. 3rd and 4th panels contain input levels of CycT1(280) and PKCε proteins (input).

**Fig. S4.**
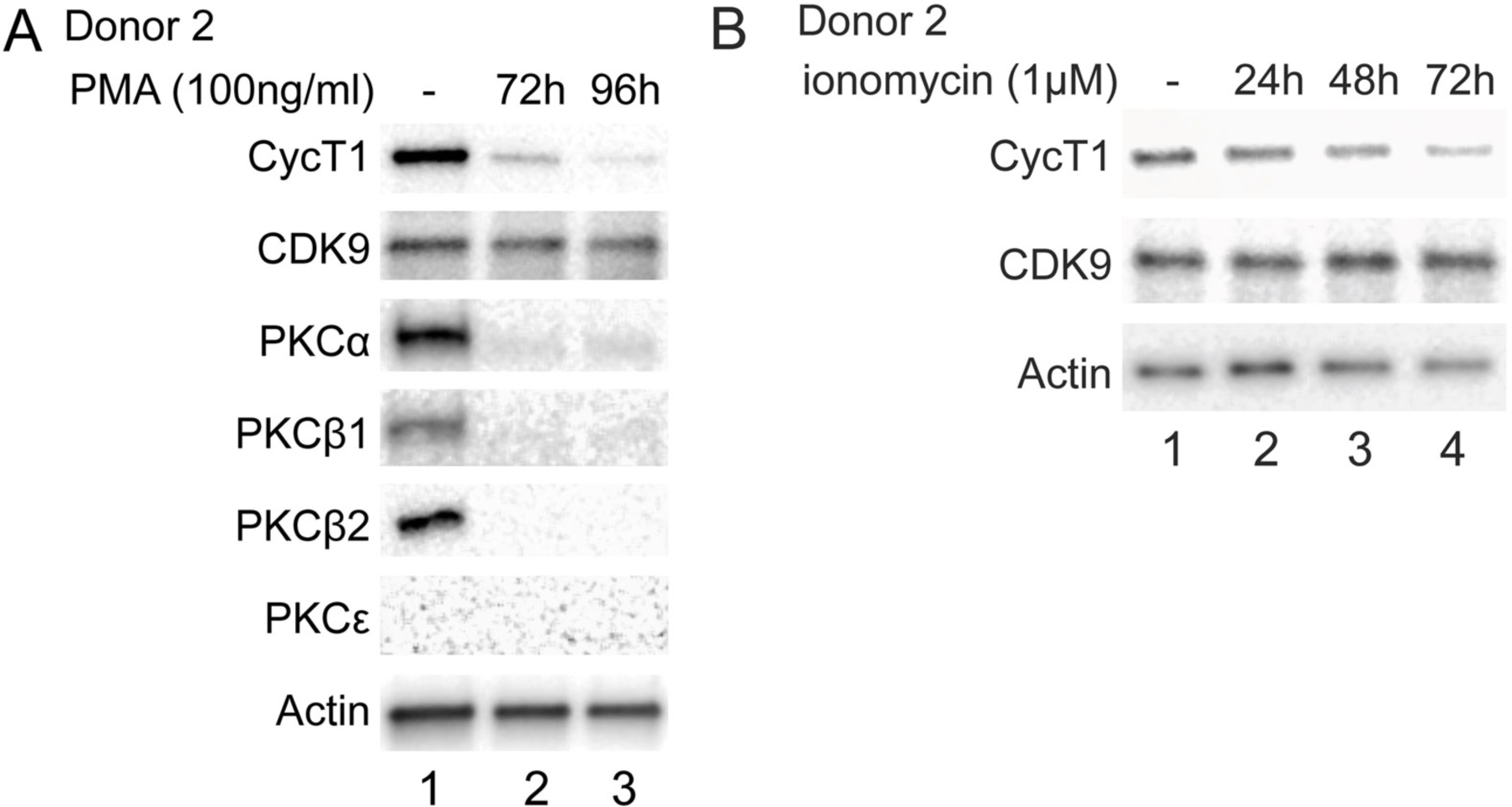
Chronic activation in primary cells decreases levels of endogenous CycT1 protein. A. Prolonged PMA treatment decreases levels of CycT1 and PKC in activated primary CD4+ T cells (donor 2). Activated primary CD4+ T cells were untreated (lane 1) or treated with 100 ng/ml PMA for 72 and 96 h (lanes 2 and 3) before cell lysis. Upper panels contain levels of endogenous CycT1, CDK9, PKCα, PKCβ1, PKCβ2, and PKCε proteins. The bottom panel contains the loading control actin protein. B. Prolonged ionomycin treatment decreases levels of CycT1 in activated primary CD4+ T cells (donor 2). Activated primary CD4+ T cells were untreated (lane 1) or treated with 1µM ionomycin for 24, 48 and 72 h (lanes 2 to 4) before cell lysis. Upper 2 panels contain levels of endogenous CycT1 and CDK9 proteins. The bottom panel contains the loading control actin protein.

## Notes

### Competing Interest Statement

The authors have declared no competing interest.

## References

AJ, C. Q., Bugai, A., & Barboric, M. (2016). Cracking the control of RNA polymerase II elongation by 7SK snRNP and P-TEFb. Nucleic Acids Res, 44(16), 7527–7539. doi:10.1093/nar/gkw585

Baumli, S., Lolli, G., Lowe, E. D., Troiani, S., Rusconi, L., Bullock, A. N., … Johnson, L. N. (2008). The structure of P-TEFb (CDK9/cyclin T1), its complex with flavopiridol and regulation by phosphorylation. EMBO J, 27(13), 1907–1918. doi:10.1038/emboj.2008.121

Cary, D. C., & Peterlin, B. M. (2020). Proteasomal Inhibition Potentiates Latent HIV Reactivation. AIDS Res Hum Retroviruses, 36(10), 800–807. doi:10.1089/AID.2020.0040

Chapman, R. D., Heidemann, M., Hintermair, C., & Eick, D. (2008). Molecular evolution of the RNA polymerase II CTD. Trends Genet, 24(6), 289–296. doi:10.1016/j.tig.2008.03.010

Cheung, T. H., & Rando, T. A. (2013). Molecular regulation of stem cell quiescence. Nat Rev Mol Cell Biol, 14(6), 329–340. doi:10.1038/nrm3591

Chiang, K., & Rice, A. P. (2012). MicroRNA-mediated restriction of HIV-1 in resting CD4+ T cells and monocytes. Viruses, 4(9), 1390–1409. doi:10.3390/v4091390

Clurman, B. E., Sheaff, R. J., Thress, K., Groudine, M., & Roberts, J. M. (1996). Turnover of cyclin E by the ubiquitin-proteasome pathway is regulated by cdk2 binding and cyclin phosphorylation. Genes Dev, 10(16), 1979–1990. doi:10.1101/gad.10.16.1979

Cohen, P. T. (2002). Protein phosphatase 1--targeted in many directions. J Cell Sci, 115(Pt 2), 241–256.

Edelson, J. R., & Brautigan, D. L. (2011). The Discodermia calyx toxin calyculin a enhances cyclin D1 phosphorylation and degradation, and arrests cell cycle progression in human breast cancer cells. Toxins (Basel), 3(1), 105–119. doi:10.3390/toxins3010105

Ferreira, M., Beullens, M., Bollen, M., & Van Eynde, A. (2019). Functions and therapeutic potential of protein phosphatase 1: Insights from mouse genetics. Biochim Biophys Acta Mol Cell Res, 1866(1), 16–30. doi:10.1016/j.bbamcr.2018.07.019

Ferrero, M., Ferragud, J., Orlando, L., Valero, L., Sanchez del Pino, M., Farras, R., & Font de Mora, J. (2011). Phosphorylation of AIB1 at mitosis is regulated by CDK1/CYCLIN B. PLoS One, 6(12), e28602. doi:10.1371/journal.pone.0028602

Franco, L. C., Morales, F., Boffo, S., & Giordano, A. (2018). CDK9: A key player in cancer and other diseases. J Cell Biochem, 119(2), 1273–1284. doi:10.1002/jcb.26293

Fujinaga, K. (2020). P-TEFb as A Promising Therapeutic Target. Molecules, 25(4). doi:10.3390/molecules25040838

Fujinaga, K., & Cary, D. C. (2020). Experimental Systems for Measuring HIV Latency and Reactivation. Viruses, 12(11). doi:10.3390/v12111279

Fujinaga, K., Irwin, D., Huang, Y., Taube, R., Kurosu, T., & Peterlin, B. M. (2004). Dynamics of human immunodeficiency virus transcription: P-TEFb phosphorylates RD and dissociates negative effectors from the transactivation response element. Mol Cell Biol, 24(2), 787–795. doi: 10.1128/mcb.24.2.787-795.2004

Garber, M. E., Wei, P., KewalRamani, V. N., Mayall, T. P., Herrmann, C. H., Rice, A. P., … Jones, K. A. (1998). The interaction between HIV-1 Tat and human cyclin T1 requires zinc and a critical cysteine residue that is not conserved in the murine CycT1 protein. Genes Dev, 12(22), 3512–3527. doi:10.1101/gad.12.22.3512

Garriga, J., Peng, J., Parreno, M., Price, D. H., Henderson, E. E., & Grana, X. (1998). Upregulation of cyclin T1/CDK9 complexes during T cell activation. Oncogene, 17(24), 3093–3102. doi:10.1038/sj.onc.1202548

Ghose, R., Liou, L. Y., Herrmann, C. H., & Rice, A. P. (2001). Induction of TAK (cyclin T1/P-TEFb) in purified resting CD4(+) T lymphocytes by combination of cytokines. J Virol, 75(23), 11336–11343. doi:10.1128/JVI.75.23.11336-11343.2001

Grana, X., De Luca, A., Sang, N., Fu, Y., Claudio, P. P., Rosenblatt, J., … Giordano, A. (1994). PITALRE, a nuclear CDC2-related protein kinase that phosphorylates the retinoblastoma protein in vitro. Proc Natl Acad Sci U S A, 91(9), 3834–3838. doi:10.1073/pnas.91.9.3834

Heissmeyer, V., Macian, F., Im, S. H., Varma, R., Feske, S., Venuprasad, K., … Rao, A. (2004). Calcineurin imposes T cell unresponsiveness through targeted proteolysis of signaling proteins. Nat Immunol, 5(3), 255–265. doi:10.1038/ni1047

Hsu, L. Y., Cheng, D. A., Chen, Y., Liang, H. E., & Weiss, A. (2017). Destabilizing the autoinhibitory conformation of Zap70 induces up-regulation of inhibitory receptors and T cell unresponsiveness. J Exp Med, 214(3), 833–849. doi:10.1084/jem.20161575

Huang, F., Shao, W., Fujinaga, K., & Peterlin, B. M. (2018). Bromodomain-containing protein 4-independent transcriptional activation by autoimmune regulator (AIRE) and NF-kappaB. J Biol Chem, 293(14), 4993–5004. doi:10.1074/jbc.RA117.001518

Ivanov, D., Kwak, Y. T., Guo, J., & Gaynor, R. B. (2000). Domains in the SPT5 protein that modulate its transcriptional regulatory properties. Mol Cell Biol, 20(9), 2970–2983. doi:10.1128/mcb.20.9.2970-2983.2000

Jadlowsky, J. K., Nojima, M., Okamoto, T., & Fujinaga, K. (2008). Dominant negative mutant cyclin T1 proteins that inhibit HIV transcription by forming a kinase inactive complex with Tat. J Gen Virol, 89(Pt 11), 2783–2787. doi:10.1099/vir.0.2008/002857-0

Jakobovits, A., Rosenthal, A., & Capon, D. J. (1990). Trans-activation of HIV-1 LTR-directed gene expression by tat requires protein kinase C. EMBO J, 9(4), 1165–1170.

Kubo, K., Ohno, S., & Suzuki, K. (1987). Primary structures of human protein kinase C beta I and beta II differ only in their C-terminal sequences. FEBS Lett, 223(1), 138–142. doi: 10.1016/0014-5793(87)80524-0

Kulkarni, A., & Bangham, C. R. M. (2018). HTLV-1: Regulating the Balance Between Proviral Latency and Reactivation. Front Microbiol, 9, 449. doi:10.3389/fmicb.2018.00449

Kuzmina, A., Verstraete, N., Galker, S., Maatook, M., Bensaude, O., & Taube, R. (2014). A single point mutation in cyclin T1 eliminates binding to Hexim1, Cdk9 and RNA but not to AFF4 and enforces repression of HIV transcription. Retrovirology, 11, 51. doi: 10.1186/1742-4690-11-51

Li, Z., Wu, J., Chavez, L., Hoh, R., Deeks, S. G., Pillai, S. K., & Zhou, Q. (2019). Reiterative Enrichment and Authentication of CRISPRi Targets (REACT) identifies the proteasome as a key contributor to HIV-1 latency. PLoS Pathog, 15(1), e1007498. doi:10.1371/journal.ppat.1007498

Lis, J. T. (2019). A 50 year history of technologies that drove discovery in eukaryotic transcription regulation. Nat Struct Mol Biol, 26(9), 777–782. doi:10.1038/s41594-019-0288-9

Lu, Z., Liu, D., Hornia, A., Devonish, W., Pagano, M., & Foster, D. A. (1998). Activation of protein kinase C triggers its ubiquitination and degradation. Mol Cell Biol, 18(2), 839–845. doi:10.1128/mcb.18.2.839

Manger, B., Weiss, A., Imboden, J., Laing, T., & Stobo, J. D. (1987). The role of protein kinase C in transmembrane signaling by the T cell antigen receptor complex. Effects of stimulation with soluble or immobilized CD3 antibodies. J Immunol, 139(8), 2755–2760.

Marshall, R. M., Salerno, D., Garriga, J., & Grana, X. (2005). Cyclin T1 expression is regulated by multiple signaling pathways and mechanisms during activation of human peripheral blood lymphocytes. J Immunol, 175(10), 6402–6411. doi:10.4049/jimmunol.175.10.6402

Mbonye, U. R., Gokulrangan, G., Datt, M., Dobrowolski, C., Cooper, M., Chance, M. R., & Karn, J. (2013). Phosphorylation of CDK9 at Ser175 enhances HIV transcription and is a marker of activated P-TEFb in CD4(+) T lymphocytes. PLoS Pathog, 9(5), e1003338. doi:10.1371/journal.ppat.1003338

Michels, A. A., & Bensaude, O. (2008). RNA-driven cyclin-dependent kinase regulation: when CDK9/cyclin T subunits of P-TEFb meet their ribonucleoprotein partners. Biotechnol J, 3(8), 1022–1032. doi:10.1002/biot.200800104

Nicoll, M. P., Proenca, J. T., & Efstathiou, S. (2012). The molecular basis of herpes simplex virus latency. FEMS Microbiol Rev, 36(3), 684–705. doi:10.1111/j.1574-6976.2011.00320.x

O’Keeffe, B., Fong, Y., Chen, D., Zhou, S., & Zhou, Q. (2000). Requirement for a kinase-specific chaperone pathway in the production of a Cdk9/cyclin T1 heterodimer responsible for P-TEFb-mediated tat stimulation of HIV-1 transcription. J Biol Chem, 275(1), 279–287. doi:10.1074/jbc.275.1.279

Palacios, E. H., & Weiss, A. (2007). Distinct roles for Syk and ZAP-70 during early thymocyte development. J Exp Med, 204(7), 1703–1715. doi:10.1084/jem.20070405

Peng, J., Zhu, Y., Milton, J. T., & Price, D. H. (1998). Identification of multiple cyclin subunits of human P-TEFb. Genes Dev, 12(5), 755–762. doi:10.1101/gad.12.5.755

Peterlin, B. M., Brogie, J. E., & Price, D. H. (2012). 7SK snRNA: a noncoding RNA that plays a major role in regulating eukaryotic transcription. Wiley Interdiscip Rev RNA, 3(1), 92–103. doi: 10.1002/wrna.106

Peterlin, B. M., & Price, D. H. (2006). Controlling the elongation phase of transcription with P-TEFb. Mol Cell, 23(3), 297–305. doi:10.1016/j.molcel.2006.06.014

Pettersen, E. F., Goddard, T. D., Huang, C. C., Couch, G. S., Greenblatt, D. M., Meng, E. C., & Ferrin, T. E. (2004). UCSF Chimera--a visualization system for exploratory research and analysis. J Comput Chem, 25(13), 1605–1612. doi:10.1002/jcc.20084

Pfeifhofer-Obermair, C., Thuille, N., & Baier, G. (2012). Involvement of distinct PKC gene products in T cell functions. Front Immunol, 3, 220. doi:10.3389/fimmu.2012.00220

Proudfoot, N. J. (2016). Transcriptional termination in mammals: Stopping the RNA polymerase II juggernaut. Science, 352(6291), aad9926. doi:10.1126/science.aad9926

Rahl, P. B., Lin, C. Y., Seila, A. C., Flynn, R. A., McCuine, S., Burge, C. B., … Young, R. A. (2010). c-Myc regulates transcriptional pause release. Cell, 141(3), 432–445. doi:10.1016/j.cell.2010.03.030

Rice, A. P. (2019). Roles of CDKs in RNA polymerase II transcription of the HIV-1 genome. Transcription, 10(2), 111–117. doi:10.1080/21541264.2018.1542254

Roche, B., Arcangioli, B., & Martienssen, R. (2017). Transcriptional reprogramming in cellular quiescence. RNA Biol, 14(7), 843–853. doi:10.1080/15476286.2017.1327510

Russo, A. A., Jeffrey, P. D., Patten, A. K., Massague, J., & Pavletich, N. P. (1996). Crystal structure of the p27Kip1 cyclin-dependent-kinase inhibitor bound to the cyclin A-Cdk2 complex. Nature, 382(6589), 325–331. doi:10.1038/382325a0

Selby, M. J., & Peterlin, B. M. (1990). Trans-activation by HIV-1 Tat via a heterologous RNA binding protein. Cell, 62(4), 769–776. doi:10.1016/0092-8674(90)90121-t

Soh, J. W., & Weinstein, I. B. (2003). Roles of specific isoforms of protein kinase C in the transcriptional control of cyclin D1 and related genes. J Biol Chem, 278(36), 34709–34716. doi: 10.1074/jbc.M302016200

Sung, T. L., & Rice, A. P. (2006). Effects of prostratin on Cyclin T1/P-TEFb function and the gene expression profile in primary resting CD4+ T cells. Retrovirology, 3, 66. doi: 10.1186/1742-4690-3-66

Sung, T. L., & Rice, A. P. (2009). miR-198 inhibits HIV-1 gene expression and replication in monocytes and its mechanism of action appears to involve repression of cyclin T1. PLoS Pathog, 5(1), e1000263. doi:10.1371/journal.ppat.1000263

Tahirov, T. H., Babayeva, N. D., Varzavand, K., Cooper, J. J., Sedore, S. C., & Price, D. H. (2010). Crystal structure of HIV-1 Tat complexed with human P-TEFb. Nature, 465(7299), 747–751. doi:10.1038/nature09131

Taube, R., Lin, X., Irwin, D., Fujinaga, K., & Peterlin, B. M. (2002). Interaction between P-TEFb and the C-terminal domain of RNA polymerase II activates transcriptional elongation from sites upstream or downstream of target genes. Mol Cell Biol, 22(1), 321–331. doi:10.1128/mcb.22.1.321-331.2002

Vorlaufer, E., & Peters, J. M. (1998). Regulation of the cyclin B degradation system by an inhibitor of mitotic proteolysis. Mol Biol Cell, 9(7), 1817–1831. doi:10.1091/mbc.9.7.1817

Wada, T., Takagi, T., Yamaguchi, Y., Ferdous, A., Imai, T., Hirose, S., … Handa, H. (1998). DSIF, a novel transcription elongation factor that regulates RNA polymerase II processivity, is composed of human Spt4 and Spt5 homologs. Genes Dev, 12(3), 343–356. doi:10.1101/gad.12.3.343

Wada, T., Takagi, T., Yamaguchi, Y., Watanabe, D., & Handa, H. (1998). Evidence that P-TEFb alleviates the negative effect of DSIF on RNA polymerase II-dependent transcription in vitro. EMBO J, 17(24), 7395–7403. doi:10.1093/emboj/17.24.7395

Wei, P., Garber, M. E., Fang, S. M., Fischer, W. H., & Jones, K. A. (1998). A novel CDK9-associated C-type cyclin interacts directly with HIV-1 Tat and mediates its high-affinity, loop-specific binding to TAR RNA. Cell, 92(4), 451–462. doi:10.1016/s0092-8674(00)80939-3

Yamaguchi, Y., Takagi, T., Wada, T., Yano, K., Furuya, A., Sugimoto, S., … Handa, H. (1999). NELF, a multisubunit complex containing RD, cooperates with DSIF to repress RNA polymerase II elongation. Cell, 97(1), 41–51. doi:10.1016/s0092-8674(00)80713-8

Zaborowska, J., Isa, N. F., & Murphy, S. (2016). P-TEFb goes viral. Bioessays, 38 *Suppl 1*, S75–85. doi:10.1002/bies.201670912

Zhou, Q., Li, T., & Price, D. H. (2012). RNA polymerase II elongation control. Annu Rev Biochem, 81, 119–143. doi:10.1146/annurev-biochem-052610-095910

